# Ultrastructural analysis of engineered rice lines reveals ferulate cross-linking as a key factor mediating lignocellulose supramolecular assembly in grass cell walls

**DOI:** 10.64898/2026.08.11.744116

**Authors:** Senri Yamamoto, Osama A. Afifi, Pingping Ji, Ryosuke Kusumi, Kayoko Kobayashi, Keisuke Kojiro, Toshiaki Umezawa, Tomoya Imai, Yuki Tobimatsu

**Author notes:** Corresponding authors: Yuki Tobimatsu and Tomoya Imai. **E-mail Address:** Senri Yamamoto, Osama A. Afifi, Pingping Ji, Ryosuke Kusumi, Kayoko Kobayashi, Keisuke Kojiro, Toshiaki Umezawa, Tomoya Imai, Yuki Tobimatsu.

## Abstract

Within the secondary cell walls of vascular plants, cellulose, hemicelluloses, and lignin associate via various covalent and non-covalent linkages to form an intricate supramolecular assembly. Although the chemical structures and cross-linking levels of the lignin– hemicellulose matrix exhibit substantial diversity *in planta*, precisely how these structural variations affect lignocellulose supramolecular assembly and macroscopic biomass properties remains largely elusive. Here, we conducted a comparative multi-scale ultrastructural analysis across engineered rice lines with targeted modifications in lignin aromatic composition and ferulate (FA)-mediated cell wall cross-linking levels. Combined solid-state nuclear magnetic resonance and wide-angle X-ray diffraction analyses revealed that depleting FA cross-linking disrupts cellulose crystalline structure and accelerates molecular mobility markedly more severely than altering the guaiacyl-to-syringyl (G/S) lignin ratio, generating a more loosened lignocellulose network. Small-angle X-ray scattering analysis further demonstrated that specific FA-depleted lines, but none of those with an altered G/S ratio, also exhibited disruptions in the nano- to mesoscale organization of cellulose microfibrils. Furthermore, FA-depleted lines generally displayed greater improvements in saccharification efficiency and more rapid thermal softening than G/S-lignin-altered lines, suggesting that disruptions in lignocellulose molecular assembly induced by FA depletion can broadly translate into macroscopic biomass properties. These findings establish a molecular basis for the pivotal role of FA cross-linking in dictating grass cell wall architecture, offering a promising structural target for advancing grass biomass utility and crop design.

## Introduction

The evolutionary innovation of the secondary cell wall endowed vascular plants with the capacity for long-distance water transport and robust structural support, driving their massive proliferation across terrestrial ecosystems. As a result, secondary cell walls, predominantly manifested as wood and non-woody fibers, constitute the most abundant form of plant-derived biomass and serve as a major reservoir of fixed carbon (Zhong et al., 2019). In the secondary cell walls, three major polymer types, i.e., cellulose, hemicelluloses, and lignin, assemble into a complex supramolecular network via both covalent and non-covalent interactions, forming a robust biocomposite termed lignocellulose (**Figure 1a**) (Terashima et al., 2009; Terrett and Dupree, 2019; Ghassemi et al., 2022). This supramolecular assembly, driven by interactions between polysaccharides and lignin, is considered a crucial determinant of cell wall biomechanical properties, thus correlating with their *in planta* functions as well as their utilization potential as biomass. While the primary structure as well as crystalline form of cellulose is highly conserved among vascular plants (Cosgrove et al., 2024), the composition and chemical structure of the other two matrix polymers, i.e., lignin and hemicelluloses, exhibit substantial diversity (Scheller and Ulvskov, 2010; Barros et al., 2015; Ralph et al., 2019; Zhong et al., 2019). These variations in the chemical structure of lignocellulose may impact its supramolecular assembly, consequently influencing downstream biomechanical properties of the cell walls as well as utilization potential as biomass. Yet, despite numerous efforts (Silveira et al., 2013; Meng and Ragauskas, 2014; Carmona *et al*., 2015; Liu et al., 2016; Shi et al., 2016; Lima et al., 2018; Martin et al., 2019; 2023; Berglund et al., 2020; Duan et al., 2021; Ménard et al., 2022; Pesquet et al., 2025; Xiao et al., 2025), the precise relationships linking the chemical structure, supramolecular organization, and physicochemical properties of lignocellulose remain poorly defined.

**Figure 1.**
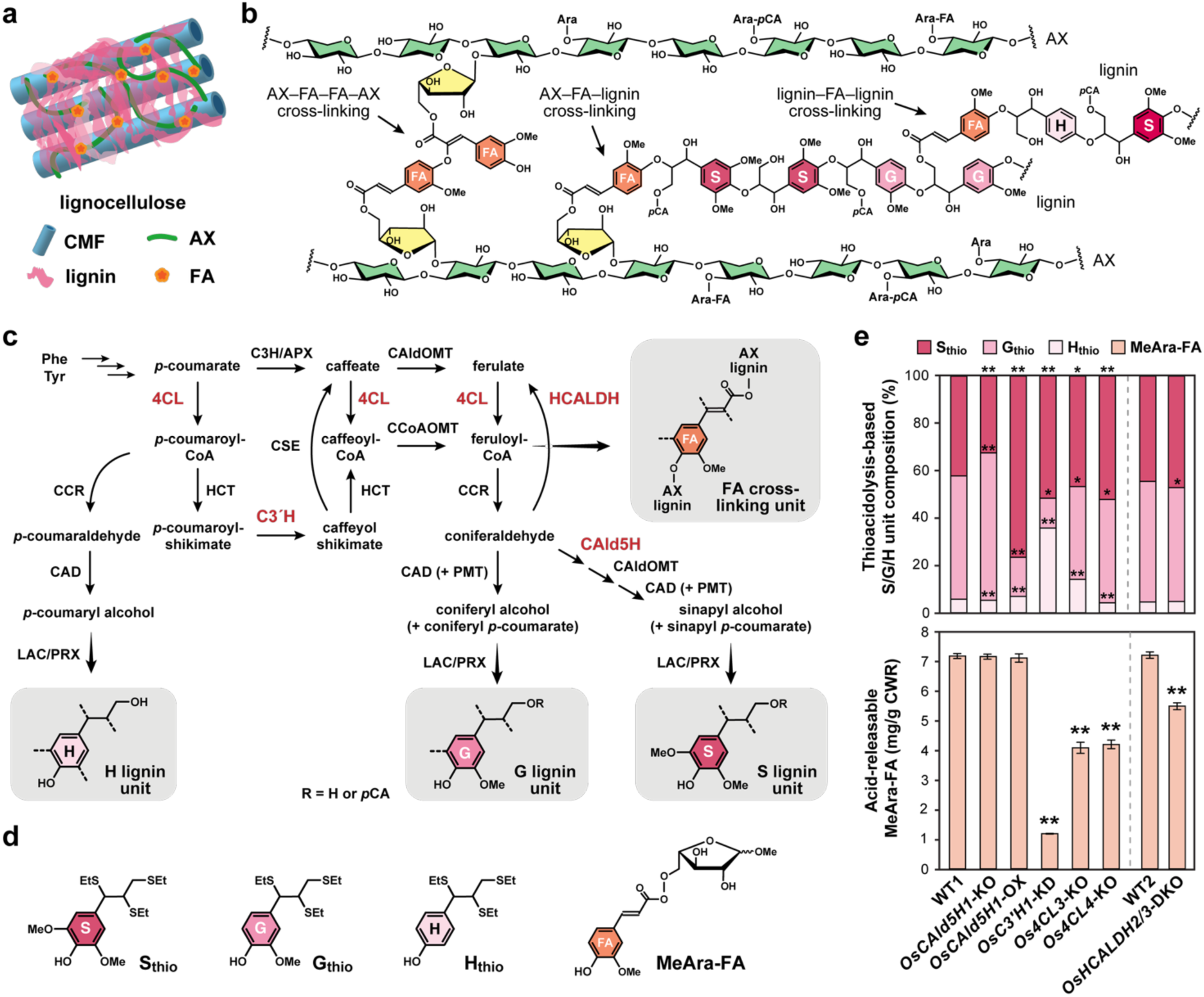
Grass lignocellulose structure and lignin- and ferulate-altered rice lines used in this study. (**a**) Illustrated model of lignocellulose organization in grass cell walls. CMF, cellulose microfibril; AX, arabinoxylan; FA, ferulate. (**b**) Chemical structures of AX and lignin, highlighting FA-mediated cross-linking. Ara, arabinose residue; *p*CA, *p*-coumarate; Me, methyl group; S: syringyl unit; G, guaiacyl unit; H, *p*-hydroxyphenyl unit. (**c**) Proposed lignin biosynthetic pathway in rice. Enzymes targeted for lignin and FA modifications in this study are highlighted in red. Phe, phenylalanine; Tyr, tyrosine; 4CL, 4-hydroxycinnamate:CoA ligase; C3H, *p*-coumarate 3-hydroxylase; APX, ascorbate peroxidase; HCT, *p*- hydroxycinnamoyl-CoA:quinate/shikimate transferase; C3′H, *p*-coumaroyl ester 3-hydroxylase; CSE, caffeoyl shikimate esterase; CAldOMT, 5-hydroxyconiferaldehyde *O*- methyltransferase; CCoAOMT, caffeoyl-CoA *O*-methyltransferase; CCR, cinnamoyl-CoA reductase; CAD, cinnamyl alcohol dehydrogenase; PMT, *p*-coumaroyl-CoA:monolignol transferase; CAld5H, coniferaldehyde 5-hydroxylase; HCALDH, hydroxycinnamaldehyde dehydrogenase;LAC, laccase; PRX, peroxidase. Monomer composition by thioacidolysis. (**d**) Lignin-derived S/G/H-type monomeric products (**S_thio_**, **G_thio_**, and **H_thio_**) released via thioacidolysis, and feruloyl arabinose (**MeAra-FA**) released via mild acid hydrolysis. (**e**) Quantification of thioacidolysis-derived lignin monomers and MeAra-FA released from culm CWR samples of the lignin-modified rice transgenic lines. Data are presented as mean ± standard deviation from three analytical replicates (*n* = 3). Asterisks (*) indicate a statistically significant difference from the respective wild-type control (Student’s *t*-test, \*\**p* < 0.01, \**p* < 0.05). CWR, cell wall residues; WT1 and WT2, wild-type control lines; *OsCAld5H1*-KO, *OsCAld5H1*-knockout line; *OsCAld5H1*-OX, *OsCAld5H1*-overexpressing line; *OsC3′H1*-KD, *OsC3′H1*-knockdown line; *Os4CL3*-KO, *Os4CL3*-knockout line; *Os4CL4*-KO, *Os4CL4*-knockout line; *OsHCALDH2/3*-DKO, *OsHCALDH2-* and *OsHCALDH3*-double-knockout line. The WT2 and *OsHCALDH2/3*-DKO lines, as well as WT1 and all other transgenic lines, were grown side-by-side.

Lignin is a complex aromatic polymer that permeates the polysaccharide matrix, embedding cellulose microfibrils (CMFs) and intimately associating with surrounding hemicellulose networks in the cell walls (**Figure 1a**). Lignin exhibits substantial structural diversity across plant lineages, cell types, and distinct cell wall layers, while also dynamically adapting to environmental cues (Barros et al., 2015; Ralph et al., 2019). Typically, gymnosperm (softwood) lignins consist predominantly of guaiacyl (G) aromatic units, whereas angiosperm lignins—encompassing basal angiosperms and eudicots (e.g., hardwoods), as well as monocots (e.g., grasses)—incorporate both G and syringyl (S) units (**Figure 1b**). In the xylem tissues of typical angiosperms, G units are enriched in vessels while S units are concentrated in fibers (Nakashima et al., 2008; Gui et al., 2020). Only small amounts of *p*- hydroxyphenyl (H) units (typically less than 5% of the total lignin) are present in normal xylem tissues across both gymnosperms and angiosperms (**Figure 1b**). However, the proportion of H units elevates markedly in specific locations such as endodermal cell corners near Casparian strips (Reyt et al., 2021) and during various biotic and abiotic stress responses (Cesarino, 2019; Kim et al., 2020). Notably, in gymnosperm compression wood formed under gravitational stress, H units can account for up to 30% of total lignin (Timell, 1986; Hiraide et al., 2021). The proportion of these major aromatic units within lignin is tightly controlled by metabolic flux through the cinnamate/monolignol pathway generating lignin monomers, including canonical monolignols (Dixon and Barros, 2019; Umezawa, 2024) (**Figure 1c**).

Distinct from cell walls developed in most of other major vascular plant lineages (gymnosperms, basal angiosperms, and eudicots), monocot grass cell walls are uniquely modified with *p*-hydroxycinnamates, mainly *p*-coumarate and ferulate (FA), bound to both lignin and hemicellulosic arabinoxylan (AX), alongside the flavone tricin, which is incorporated specifically into lignin (Ralph et al., 2010, 2019). Among these unique cell wall modifications, FA serves as a critical cross-linker between lignin and AX (**Figure 1b**). The majority of this FA is esterified to the *O*-5 position of arabinosyl side chains on AX (Ralph, 2010), while a minor fraction of FA is linked to the γ-position of monolignols, leading to the production of γ-feruloylated lignin polymer units upon cell wall lignification (Karlen et al., 2016). During cell wall maturation, these AX- and monolignol-bound FAs participate in oxidative radical coupling with one another, as well as with free lignin monomers and growing lignin polymers, generating an extensive network that highly cross-link the cell wall matrix (**Figure 1b**) (Ralph, 2010; Karlen et al., 2016; Ralph et al., 2019). The establishment of such FA-mediated cross-linking network is hypothesized to play a pivotal role not only in the physiological functions of grass cell walls but also in dictating the utilization properties of grass biomass (Buanafina, 2009; Ralph, 2010; de Oliveira et al., 2015; Hatfield et al., 2017; de Souza et al., 2018; Mnich et al., 2020; Chandrakanth et al., 2023; Smith and Ralph, 2024; Yamamoto et al., 2024; Yang et al., 2004).

Such substantial diversity of lignin structure and cell wall cross-linking underscores the complex biophysical mechanisms governing cell wall architecture. However, the mechanistic links translating these biochemical variations into the supramolecular assembly and macroscopic properties of lignocellulose remain elusive. Here, we aimed to close this gap by performing comparative ultrastructural analyses on a series of mutant and transgenic rice lines with targeted modifications in lignin aromatic composition and FA-mediated cell wall cross-linking levels. Combined wide-angle X-ray diffraction (WAXD) and solid-state NMR (ssNMR) analyses resolved cellulose crystallinity and polymer mobility within the cell wall matrix. In parallel, small-angle X-ray scattering (SAXS) characterized nano- to mesoscale cell wall architecture, focusing on CMF organization. Finally, enzymatic saccharification efficiency and thermal softening behavior were comparatively evaluated to correlate the structural shifts of lignocellulose with biomass properties.

## Results

### Rice lines with altered lignin composition and cell wall cross-linking FA levels

For a comparative analysis of lignocellulose supramolecular assembly and biomass properties, we utilized a series of rice transgenic and mutant lines with altered lignin aromatic composition and cell wall-bound FA levels. These rice lines, generated via targeted down- or over- expression of the cinnamate/monolignol pathway enzymes (**Figure 1c**), include the *CAld5H*-knock-out line (*OsCAld5H1*-KO) (Takeda et al., 2019a), the *CAld5H*-over-expression line (*OsCAld5H1*-OX) (Takeda et al., 2017), the *C3ʹH*-knock-down line (*OsC3ʹH1*-KD) (Takeda et al., 2018), the *4CL*-knock-out lines (*Os4CL3*-KO and *Os4CL4*-KO) (Afifi et al., 2022), and the *HCALDH*-double-knock-out line (*OsHCALDH2/3*-DKO) (Yamamoto et al., 2024). As confirmed below, *OsCAld5H1*-KO, *OsCAld5H1*-OX, and *OsC3ʹH1*-KD represent lignin aromatic composition-altered lines with enrichments in G, S and H units, respectively, while *Os4CL3*-KO, *Os4CL4*-KO, and *OsHCALDH2/3*-DKO lines represent cell wall-bound FA-depleted lines with minor changes in lignin composition. Notably, *OsC3ʹH1*-KD has been reported to exhibit depleted cell wall-bound FA levels along with H lignin enrichment (Takeda et al., 2018). The rice transgenic and mutant lines were grown to maturity alongside their respective wild-type (WT) controls; WT1 served as the control for all lines except *OsHCALDH2/3*-DKO, for which WT2 was used (**Table S1**).

### Lignocellulose composition analyses of cell wall-altered rice lines

Prior to analyzing lignocellulose supramolecular structure, we confirmed the cell wall chemotypes of the rice mutant and transgenic lines by subjecting extractive-free cell wall residue (CWR) samples from mature culms to a series of chemical methods. Alterations in lignin aromatic composition were verified by analytical thioacidolysis, which quantifies lignin-derived H-, G-, and S-type compounds (**S_thio_**, **G_thio_**, and **H_thio_**) released via chemical cleavage of dominant β–O–4 ether linkages (**Figure 1d**). Consistent with previous reports (Takeda et al., 2017; 2018; 2019a), the relative abundance of G-type compounds significantly increased in *OsCAld5H1*-KO (by ∼17%), S-type compounds increased in *OsCAld5H1*-OX (by ∼85%), and H-type compounds increased in *OsC3′H1*-KD (by ∼500%) compared to the corresponding WT control (**Figure 1e**). In contrast, the S/G/H compound ratio remained largely unchanged in the *Os4CL3*-KO, *Os4CL4*-KO, and *OsHCALDH2/3*-DKO lines relative to the corresponding WT controls (**Figure 1e**), which overall aligns with previous findings (Afifi et al., 2022; Yamamoto et al., 2024). The extent of FA-mediated cell wall cross-linking was evaluated by quantifying free FA released via mild alkaline hydrolysis and feruloylated arabinose (methyl 5-*O*-feruloyl-L-arabinose, **MeAra-FA**) (**Figure 1d**) released via methanolic acidolysis of CWRs (Lapierre et al., 2018; Yamamoto et al., 2024). Compared with WT controls, the alkaline-releasable FA content was significantly decreased in *OsC3′H1*-KD (by ∼77%), *Os4CL3*-KO (by ∼33%), *Os4CL4*-KO (by ∼38%), and *OsHCALDH2/3*-DKO (by ∼22%), whereas *OsCAld5H1*-KO and *OsCAld5H1*-OX showed no significant change (**Table 1**), which aligns with previous reports (Takeda et al., 2017, 2018, 2019a; Afifi et al., 2022; Yamamoto et al., 2024). Similarly, significant reductions in **MeAra-FA** released via methanolic acidolysis were detected in *OsC3′H1*-KD (by ∼83%), *Os4CL3*-KO (by ∼43%), *Os4CL4*-KO (by ∼42%), and *OsHCALDH2/3*-DKO (by ∼25%), whereas *OsCAld5H1*-KO and *OsCAld5H1*-OX showed no significant changes relative to WT controls (**Figure 1c**). Overall, these analyses verified the expected modifications to lignin aromatic composition and cell wall-bound FA levels within each respective line.

**Table 1.** Lignin, ferulate, and polysaccharide composition in culm cell walls from lignin- and ferulate-altered rice lines.

| Component<br>(mg/g CWR) | WT1 | *OsCald5H1*-KO | *OsCald5H1*-OX | *OsC3'H1*-KD | *Os4CL3*-KO | *Os4CL4*-KO | WT2 | *OsHCALDH2/3*-DKO |
| --- | --- | --- | --- | --- | --- | --- | --- | --- |
| Klason lignin | 140.6 ± 1.5 | 145.7 ± 2.8 | **115.9 ± 3.6\*\*** | **102.6 ± 9.3\*\*** | **88.0 ± 3.2\*\*** | **117.1 ± 0.8\*\*** | 130.2 ± 1.0 | **121.4 ± 2.5\*\*** |
| Cell wall-bound<br>ferulate | 5.1 ± 0.07 | 4.9 ± 0.10 | 4.9 ± 0.15 | **1.2 ± 0.02\*\*** | **3.4 ± 0.20\*\*** | **3.2 ± 0.01\*\*** | 5.6 ± 0.14 | **4.4 ± 0.04\*\*** |
| Neutral sugars |  |  |  |  |  |  |  |  |
| Crystalline glucose<sup>a</sup> | 428.4 ± 15.7 | 409.9 ± 18.2 | **375.7 ± 10.1\*\*** | 422.0 ± 0.5 | **397.2 ± 6.1\*** | **369.6 ± 9.8\*\*** | 366.9 ± 4.2 | 357.3 ± 19.5 |
| Amorphous glucose<sup>b</sup> | 28.4 ± 1.2 | 26.1 ± 2.0 | **41.2 ± 0.8\*\*** | **35.7 ± 0.2\*\*** | **48.0 ± 0.7\*\*** | **41.6 ± 3.4\*\*** | 30.8 ± 0.4 | **36.4 ± 2.1\*\*** |
| Xylose | 75.2 ± 1.6 | 72.0 ± 5.4 | 76.5 ± 1.5 | 73.9 ± 0.8 | **80.9 ± 0.8\*\*** | 75.7 ± 4.9 | 79.1 ± 0.9 | 80.7 ± 4.2 |
| Arabinose | 22.1 ± 0.6 | 20.8 ± 1.6 | 22.4 ± 0.8 | 23.5 ± 0.5 | **25.6 ± 0.8\*\*** | **26.4 ± 1.5\*\*** | 25.1 ± 0.3 | 25.7 ± 1.3 |
| Galactose | 14.2 ± 0.5 | 13.1 ± 0.7 | 14.6 ± 0.3 | 14.4 ± 0.2 | **17.9 ± 0.3\*\*** | **20.3 ± 1.5\*\*** | 15.7 ± 0.3 | 16.6 ± 0.6 |

<sup>1</sup>Glucose released from trifluoroacetic acid-insoluble cell wall fractions (glucose primarily from crystalline cellulose). <sup>2</sup>Glucose released from trifluoroacetic acid-soluble cell wall fractions (glucose primarily from amorphous cellulose and hemicelluloses). Data are presented as mean ± standard deviation from three analytical replicates (*n* = 3). Asterisks (\*) and numbers in bold indicate a statistically significant difference from the respective wild-type control (Student's *t*-test, \*\**p* < 0.01, \**p* < 0.05). Values highlighted in bold indicate significant differences from the corresponding WT controls. CWR, cell wall residues; WT1 and WT2, wild-type control lines; *OsCald5H1*-KO, *OsCald5H1*-knockout line; *OsCald5H1*-OX, *OsCald5H1*-overexpressing line; *OsC3'H1*-KD, *OsC3'H1*-knockdown line; *Os4CL3*-KO, *Os4CL3*-knockout line; *Os4CL4*-KO, *Os4CL4*-knockout line; *OsHCALDH2/3*-DKO, *OsHCALDH2*- and *OsHCALDH3*-double-knockout line. The WT2 and *OsHCALDH2/3*-DKO lines, as well as WT1 and all other transgenic lines, were grown side-by-side.

As an essential baseline prior to analyzing lignocellulose supramolecular assembly, we also evaluated overall lignin and sugar compositions via Klason and neutral sugar assays. Klason lignin content was moderately decreased (7–37%) in all rice lines except *OsCAld5H1*- KO relative to WT controls (**Table 1**); thus, these differences in lignin content must be accounted for when interpreting subsequent supramolecular variations among these lines, as discussed below. Neutral sugar analysis showed that glucose released from the trifluoroacetic acid (TFA)-insoluble cell wall fraction, representing crystalline cellulose, significantly decreased in *OsCAld5H1*-OX, *Os4CL3*-KO, and *Os4CL4*-KO, but remained unchanged in *OsCAld5H1*-KO, *OsC3′H1*-KD, and *OsHCALDH2/3*-DKO (**Table 1**). Conversely, glucose released from the TFA-soluble cell wall fraction, presumed to originate from amorphous cellulose and hemicellulosic glucans, significantly increased in all lines except *OsCAld5H1*- KO (**Table 1**). Furthermore, xylose, arabinose, and galactose released from hemicelluloses significantly increased in *Os4CL3*-KO and *Os4CL4*-KO (except for xylose) compared to WT, while remaining unaffected in all other lines (**Table 1**). These shifts suggest proportional increase of cellulose and hemicelluloses against reduced lignin as well as underlying modifications to cellulose crystalline structure, which we further examined below.

### Analysis of cellulose crystalline structure by WAXD and ssNMR

As a first step in evaluating lignocellulose assembly in the lignin- and FA-modified rice cell walls, we investigated cellulose crystalline structure using WAXD and ssNMR.

The WAXD profiles of culm CWR samples were subjected to multi-peak fitting to resolve crystalline cellulose reflections from amorphous contributions (which encompass amorphous cellulose as well as the hemicellulose and lignin matrix) (**Figure 2a; Figure S1**). Apparent crystallinity index (*CrI*) calculations using the Segal method (Segal et al., 1959) revealed that lignocellulose crystallinity significantly decreased in *OsCAld5H1*-OX (by ∼14%), *OsC3′H1*-KD (by ∼12%), *Os4CL3*-KO (by ∼12%), *Os4CL4*-KO (by ∼45%), and *OsHCALDH2/3*-DKO (by ∼22%) compared with WT controls, whereas no significant change was observed in *OsCAld5H1*-KO (**Figure 2b**). In contrast, cellulose crystal size, as determined by the Scherrer equation (Scherrer, 1918), remained unaffected across all the lignin- and FA-modified lines examined, although a slight reduction was observed in *OsHCALDH2/3*-DKO relative to the WT2 control (**Figure 2c**).

**Figure 2.**
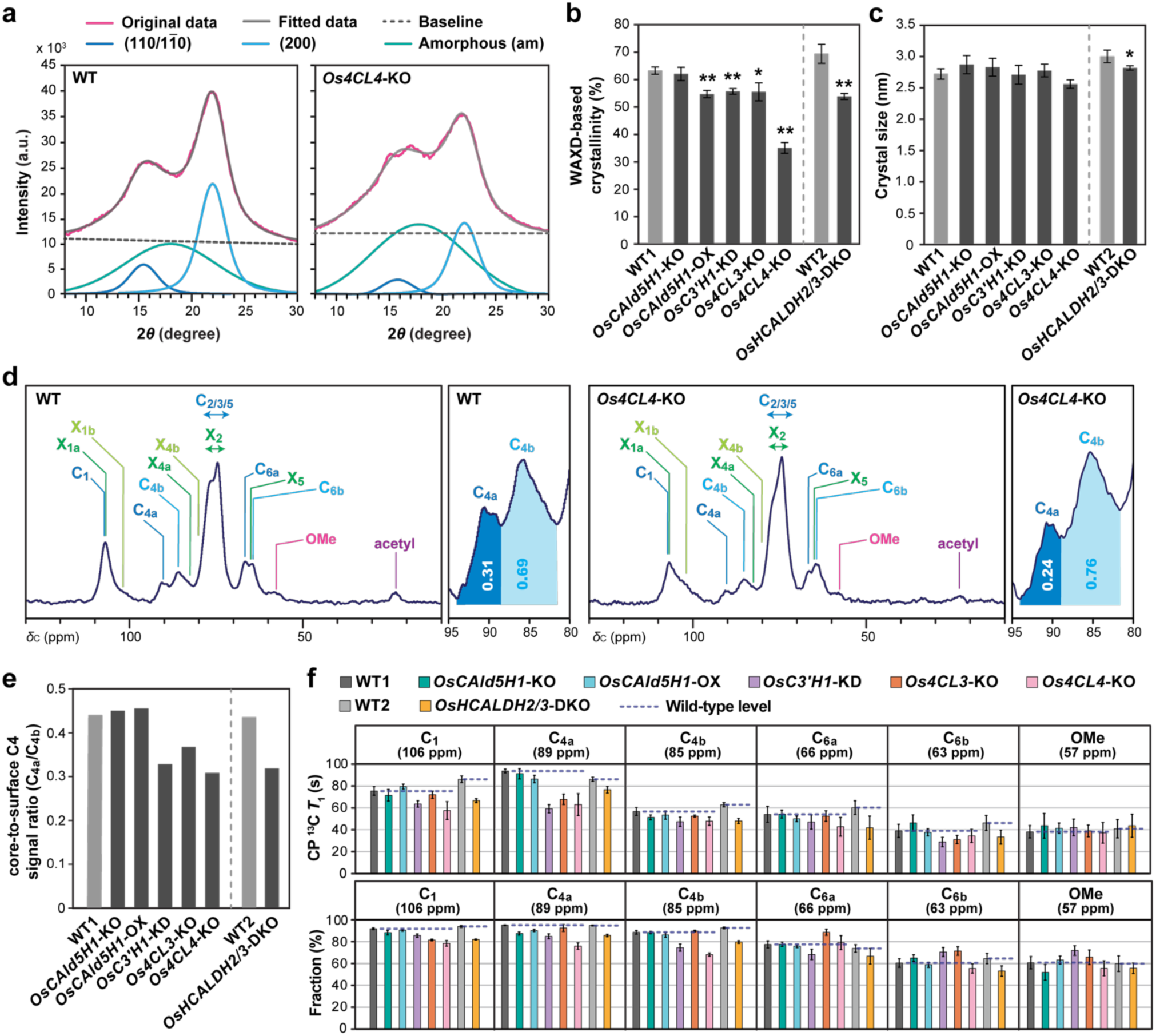
WAXD and ssNMR analysis of lignin- and ferulate-altered rice cell walls. (**a**) Representative wide-angle X-ray diffraction (WAXD) profiles of culm cell walls from WT1 and *Os4CL4*-KO plants. The experimental WAXD profiles (red line) were deconvoluted via multi-peak fitting (gray line) into an amorphous contribution (green line) and crystalline cellulose reflections corresponding to the (200) and (110/1 1 0) lattice planes (light and dark blue lines). The dashed gray line indicates the baseline subtracted prior to fitting. Additional profiles for all other lines are provided in **Figure S1**. (**b** and **c**) WAXD-derived cell wall crystallinity (**b**) and crystal size (**d**) data determined using the Segal method and Scherrer equation, respectively. The values are means ± standard deviation from three analytical runs (*n* = 3). Asterisks (*) indicate a statistically significant difference from the respective wild-type control (Student’s *t*-test, \*\**p* < 0.01, \**p* < 0.05). (**d**) Representative ^13^C CP-MAS solid-state NMR (ssNMR) spectra of culm cell walls from WT and *Os4CL4*-KO plants. Peak assignments and additional profiles for all other lines are provided in **Table S2** and **Figure S2**. (**e**) Cellulose core-to-surface C4 signal ratio (**C_4a_**/**C_4b_**). (**f**) CP ^13^C spin-lattice relaxation time (*T*_1_) data for major cellulose and lignin methoxy (**OMe**) carbon sites. Delay time-dependent signal decays were fitted to a double exponential function to extract independent *T*_1_ values for slow- and fast-relaxing components. Top panels display *T*_1_ values, and bottom panels display fractional abundances (normalized to slow + fast = 100%), for the slow-relaxing components. Complete datasets, including all fast-relaxing component parameters, are listed in **Table S3**. Error bars represent the standard deviation of the curve-fitting coefficients. WT1 and WT2, wild-type control lines; *OsCAld5H1*-KO, *OsCAld5H1*-knockout line; *OsCAld5H1*-OX, *OsCAld5H1*-overexpressing line; *OsC3′H1*-KD, *OsC3′H1*-knockdown line; *Os4CL3*-KO, *Os4CL3*-knockout line; *Os4CL4*-KO, *Os4CL4*-knockout line; *OsHCALDH2/3*-DKO, *OsHCALDH2-* and *OsHCALDH3*-double-knockout line. The WT2 and *OsHCALDH2/3*-DKO lines, as well as WT1 and all other transgenic lines, were grown side-by-side.

To further probe cellulose crystallinity, ^13^C magic-angle spinning (MAS) ssNMR spectra were collected using ^1^H–^13^C cross-polarization (CP) to selectively enhance signals from rigid cellulose domains over those of more mobile hemicelluloses and lignin domains. Accordingly, all CP-MAS spectra of the rice cell wall samples were dominated by cellulose signals originating from glucose residues in two distinct environments, which often assigned as “internal/crystalline” and “surface/noncrystalline” cellulose domains (Simmons et al., 2016) (**Figure 2d; Figure S2; Table S2**). Although the precise structural origins of these separated signals remain under active investigation (Cresswell et al., 2025), we utilized the traditional C4 signal ratio (**C_4a_** at 89 ppm for internal/crystalline cellulose; **C_4b_** at 85 ppm for surface/noncrystalline cellulose) as a practical metric to assess relative differences in the cellulose crystallinity or the internal-versus-surface molecular environments of CMFs among the mutant lines (**Figure 2d; Figure S2; Table S2**). Notably, the **C_4a_**/**C_4b_** ratio significantly decreased in all the FA-depleted lines, i.e., *OsC3′H1*-KD (by ∼19%), *Os4CL3*-KO (by ∼11%), *Os4CL4*-KO (by ∼23%), and *OsHCALDH2/3*-DKO (by ∼22%) (**Figure 2e**). In contrast, the **C_4a_**/**C_4b_** ratio remained unchanged in the two lines with altered G/S lignin composition, i.e., *OsCAld5H1*-KO and *OsCAld5H1*-OX (**Figure 2e**).

Overall, the WAXD- and ssNMR-derived crystallinity measurements align well across all lines except *OsCAld5H1*-OX, where apparent macro-scale lignocellulose crystallinity decreased (**Figure 2b**) despite unaltered molecular-level cellulose crystallinity (**Figure 2e**). This discrepancy likely reflects increased amorphous background X-ray scattering from the non-cellulosic polymer matrix rather than a true structural modification of the cellulose lattice. Taken together, these findings demonstrate that reduced FA-mediated cell wall cross-linking contributes to disrupted cellulose crystalline structure, whereas alterations in lignin aromatic composition—particularly shifts in the major G/S monomeric ratio—have minimal impact on cellulose crystallinity.

### Analysis of lignocellulose molecular mobility by ssNMR

We also assessed the molecular mobility of cellulose and lignin components in the lignin- and FA-modified rice cell walls based on site-specific ^13^C spin-lattice relaxation time (*T*_1_) data, which are sensitive to nanosecond-timescale motions of cell wall polymers (García et al., 2011; Wang et al., 2014; Kang et al., 2019; Martin et al., 2019, 2023). We employed a Torchia CP pulse sequence (Torchia, 1978) to collect *T*_1_ data for cellulose ring carbons (**C_1_**–**C_6a/b_**) as well as the lignin methoxy carbon (**OMe**). Consistent with earlier cell wall studies using analogous ssNMR relaxation approaches (Horii et al., 1984; Zuckerstätter et al., 2013; Martin et al., 2019, 2023), the obtained magnetization decay curves were well fitted by a double-exponential function, revealing two distinct relaxing pools for each carbon site: a relatively slower-relaxing (rigid) fraction and a faster-relaxing (mobile) fraction.

Compared with WT controls, all four FA-depleted lines (*OsC3′H1*-KD, *Os4CL3*-KO, *Os4CL4*-KO, and *OsHCALDH2/3*-DKO) displayed notably decreased *T*_1_ values across most cellulose carbon sites (**Figure 2f; Table S3**). The fractional weighting of the slower-relaxing component at these cellulose carbon sites was generally similar to or lower than that of the WT controls (**Figure 2f; Table S3**). In contrast, neither cellulose *T*_1_ values nor fractional weightings were affected in the two lignin-modified lines, *OsCAld5H1*-KO and *OsCAld5H1*- OX (**Figure 2f; Table S3**). These results indicate that, while alterations in G/S lignin composition exert negligible impact on cellulose dynamics, the molecular mobility of cellulose is markedly increased in rice cell walls with reduced FA cross-linking, reflecting alleviated intermolecular constraints and a loosened lignocellulose network. In contrast, no major changes were observed in *T*_1_ values or fractional weightings for the lignin methoxy carbon in any of the examined samples, suggesting that, at least within the timescale and nano-domain sizes probed by the current CP-*T*_1_ measurements, overall lignin molecular mobility was not perturbed in any of the modified lines (**Figure 2f; Table S3**).

### Analysis of CMF organization by SAXS

To further investigate the lignocellulose supramolecular assembly in lignin- and FA-altered rice cell walls, we analyzed intact rice culm samples using SAXS, a robust technique for probing nano-to-mesoscale organization of lignocellulose within plant cell walls (Nishiyama, 2009; Xu et al., 2013; Nishiyama et al., 2014; Martínez-Sanz et al., 2015; Penttilä et al., 2019; 2020; Horiyama et al., 2022; Hua et al., 2024; Hu et al., 2025; 2026; Tanaka et al., 2025). Synchrotron X-rays were directed radially through dry-state rice culms from the inner to the outer wall surface; due to high beam penetration, the resulting data represent a superposition of scattering signals across the entire illuminated wall thickness. The collected SAXS patterns were highly anisotropic, reflecting the preferential alignment of CMFs in the secondary cell walls (**Figure S3**). The azimuthal region of maximum scattering intensity (the equatorial direction) reflects the lateral packing density and nanoscale spatial arrangement of CMFs within the secondary cell walls (Nishiyama, 2009; Penttilä et al., 2019).

To evaluate CMF orientation across the rice cell wall samples, we generated azimuthal scattering intensity profiles (*χ*–*I* plots) within the scattering vector range *q* = 0.075–0.25 Å⁻¹ and performed multi-peak fitting. Following Lichtenegger et al. (1999), we deconvolved the signals into a strong central peak flanked by two peaks of comparable intensity, accounting for the geometric superposition of X-ray scattering contributions from the radial and tangential cell walls, respectively (**Figure 3a; Figure S4**). The full width at half maximum (FWHM) of the central radial peak, which represents the overall orientational heterogeneity of CMFs (Lichtenegger et al. 1999), remained generally consistent across most of the examined rice lines, but markedly increased in *Os4CL4*-KO compared with its WT control (WT1), indicating that CMF orientation is significantly disordered in the cell walls of this particular mutant (**Figure 3b**). Next, equatorial scattering intensity (*I* vs. scattering vector *q*; *q*–*I* plots) profiles were analyzed to further examine the nanoscale arrangement of CMFs within the lignin– hemicellulose matrix (**Figure 3c; Figure S5**). A distinct shoulder peak around *q* = 0.1–0.2 Å⁻¹, attributed to the electron density contrast between the CMFs and the surrounding cell wall matrix (Nishiyama, 2009; Penttilä et al., 2019), was clearly observed in WT controls and most mutant and transgenic culm samples (**Figure 3c; Figure S5**). Notably, this shoulder peak was substantially diminished in *Os4CL4*-KO, suggesting a decreased electron density contrast between the CMFs and the matrix. This change was further corroborated by Kratky plots (*q*²*I* vs. *q*), where the prominent peak maxima at *q* = 0.1–0.2 Å⁻¹ present in WT controls and other modified lines were markedly suppressed in *Os4CL4*-KO (**Figure 3c; Figure S6**). Furthermore, *OsC3′H1*-KD displayed notably elevated scattering intensity around *q* = 0.05 Å⁻¹ relative to WT controls and all other lines, as clearly indicated in both *q*–I and Kratky plots (**Figure 3c; Figure S5; Figure S6**). Given that scattering at *q* = 0.1–0.2 Å^−1^ is attributed to individual CMFs, the scattering signal at *q* ≈ 0.05 Å^−1^ corresponds to larger real-space dimensions. This indicates that *OsC3′H1*-KD undergoes higher-order structural alterations at a length scale exceeding that of individual CMFs, such as microfibril aggregation or bundling (Penttilä et al., 2019, 2020).

**Figure 3.**
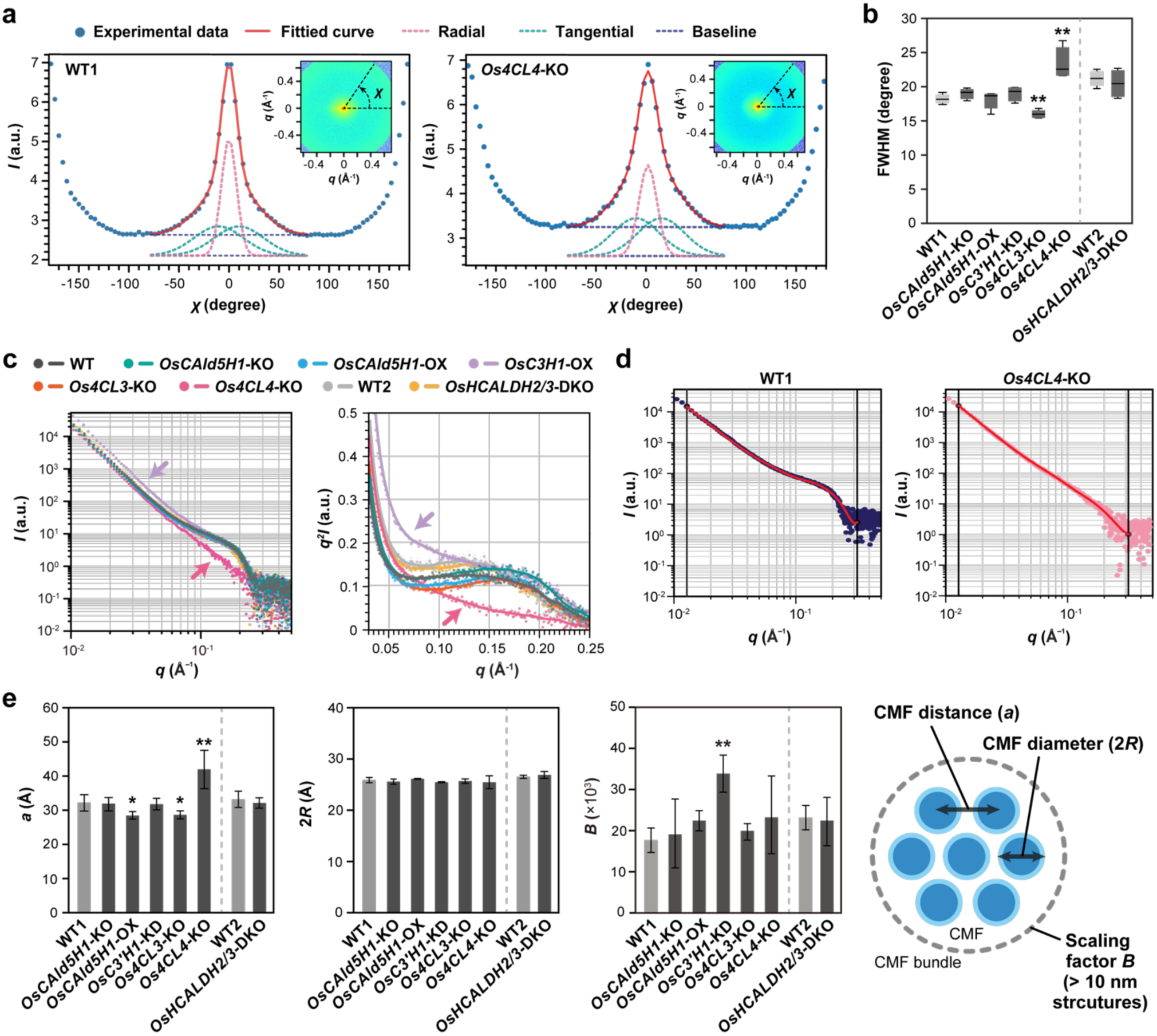
SAXS analysis of lignin- and ferulate-altered rice cell walls. (**a**) Representative small-angle X-ray scattering (SAXS) profile (inset) and azimuthal intensity profile (*χ-I* plot) of culm cell walls from WT1 and *Os4CL4*-KO plants. The *χ-I* plot was fitted with three Gaussian functions across the range *q* = 0.075-0.25 Å^−1^ f to resolve one radial and two tangential orientations. Additional profiles across all lines are provided in **Figures S3** and **S4**. (**b**) Full width at half maximum (FWHM) of the radial Gaussian fit. (**c**) Representative background-subtracted equatorial *q-I* and Kratky plots of culm cell walls from lignin-and ferulate-altered rice lines. Arrows indicate increased and decreased scattering features in *OsC3′H1*-KD and *Os4CL4*-KO lines. Additional profiles obtained from four biologically independent rice culm samples are provided in **Figures S5** and **S6**. (**d** and **e**) SAXS data fitting analysis using the WoodSAS model (Penttilä et al., 2019). Representative fitting results of equatorial *q*–*I* plots for culm cell walls from WT1 and *Os4CL4*-KO plants (**d**) and the resulting cellulose microfibril (CMF) packing parameters across all lines (**e**) are shown. In **e**, the WoodSAS-derived CMF distance (*a*), diameter (2*R*), and constant *B* term of the model equation are reported. The complete dataset is provided in **Table S5**. In **b** and **e**, data represent means ± standard deviation from four biologically independent plants (*n* = 4). Asterisks (*) indicate a statistically significant difference from the respective wild-type control (Student’s *t*- test, \*\**p* < 0.01, \**p* < 0.05). WT1 and WT2, wild-type control lines; *OsCAld5H1*-KO, *OsCAld5H1*-knockout line; *OsCAld5H1*-OX, *OsCAld5H1*-overexpressing line; *OsC3′H1*-KD, *OsC3′H1*-knockdown line; *Os4CL3*-KO, *Os4CL3*-knockout line; *Os4CL4*-KO, *Os4CL4*-knockout line; *OsHCALDH2/3*-DKO, *OsHCALDH2-* and *OsHCALDH3*-double-knockout line. WT2 and *OsHCALDH2/3*-DKO, and WT1 and all other rice lines, were grown side-by-side.

To interpret the SAXS data more quantitatively, we analyzed the equatorial scattering profiles (*q–I* plots) by subjecting them to multi-parameter fitting using the WoodSAS model, which assumes individual CMFs as infinite cylinders packed in a hexagonal array with paracrystalline distortion (**Figure 3d**) (Penttilä et al., 2019). While the WoodSAS model was initially designed to characterize CMF organization in wood cell walls, it has recently been applied to analyze SAXS data from grass cell walls (Hua et al., 2024; Hu et al., 2026). The best-fit parameters obtained for our rice culm cell walls (**Table S4**) were generally consistent with those previously determined for other cell wall samples using identical methodologies (Penttilä et al., 2019; 2020; Horiyama et al., 2022; Hua et al., 2024; Tanaka et al., 2025; Hu et al., 2026). Notably, the interfibrillar distance *a* was significantly increased in *Os4CL4*-KO, rising from approximately 3.3 nm in the WT1 control to approximately 4.1 nm, whereas no marked changes were observed for any of the other modified lines (**Figure 3e; Table S4**). Most remaining WoodSAS fitting parameters, including the CMF diameter *2R*, exhibited no significant alterations across any of the mutant or transgenic lines relative to WT controls (**Figure 3e; Table S4**). However, the second scaling factor (*B*), which primarily reflects scattering contributions in the *q*-range of 0.01–0.1 Å⁻¹ and thus accounts for structural features larger than 10 nm (i.e., length scales exceeding individual CMFs) (Penttilä et al., 2019; 2020), was significantly elevated in the *OsC3′H1*-KD line compared with the WT controls and all other modified lines (**Figure 3e; Table S4**). Thus, consistent with our qualitative evaluations based on the *q–I* and Kratky profiles, quantitative analysis using the WoodSAS model confirms that CMF spatial organization is substantially disrupted in the two FA-depleted mutants, *Os4CL4*-KO and *OsC3′H1*-KD, whereas CMF organization remained largely intact across the remaining lignin- and FA-altered lines.

### Enzymatic saccharification and thermal softening

Finally, to link the supramolecular structural changes of lignocellulose with macroscopic biomass properties, we compared the enzymatic saccharification efficiency and thermal softening behavior of rice cell wall samples from the lignin- and FA-altered lines against WT controls.

#### Enzymatic saccharification

Destarched culm CWRs were subjected to digestion with a cellulolytic enzyme cocktail both with and without mild alkaline pretreatment (Takeda et al., 2019b; Yamamoto et al., 2024). Without pretreatment, all lines with reduced FA cross-linking, except *Os4CL4*-KO, exhibited significantly enhanced saccharification efficiency, whereas the two lines with altered G/S lignin composition, i.e., *OsCAld5H1*-KO and *OsCAld5H1*-OX, displayed saccharification efficiency similar to the WT controls (**Figure 4a**). These results are fully consistent with previous non-pretreated saccharification assays conducted for each lignin- and FA-altered line (Takeda et al., 2018, 2019b; Afifi et al., 2022; Yamamoto et al., 2024). Following alkaline pretreatment, both lignin- and FA-altered lines showed substantial improvements in saccharification efficiency, though gains were overall higher in the FA-depleted mutants compared to the G/S-lignin-altered lines (**Figure 4a**).

**Figure 4.**
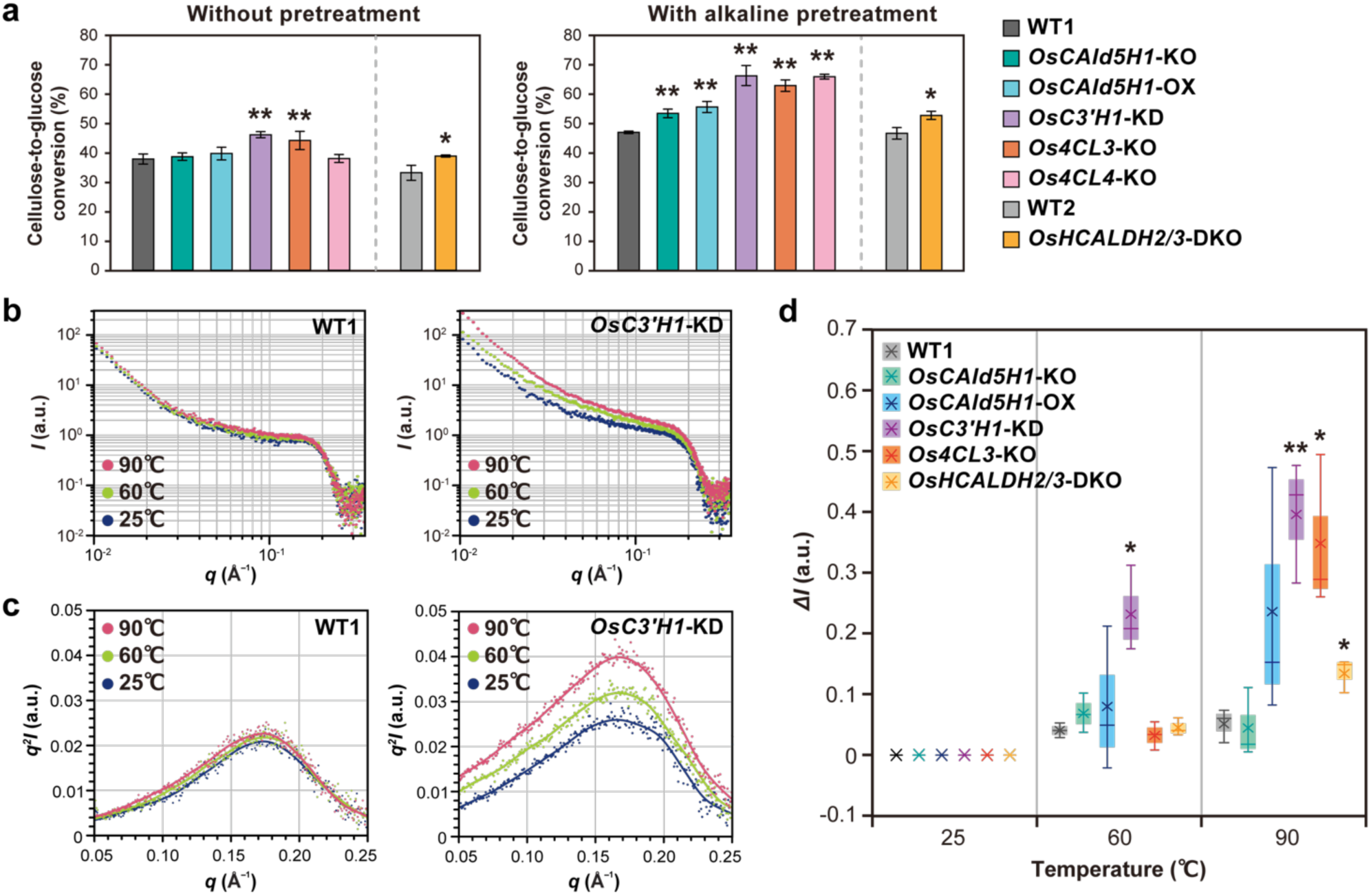
Enzymatic saccharification and thermal softening of lignin- and ferulate-altered rice cell walls. (**a**) Cellulose-to-glucose conversion rate following enzymatic saccharification of unpretreated and alkaline-pretreated rice culm cell walls. Assays were conducted on pooled cell wall residue (CWR) samples prepared from three biologically independent plants per line. (**b**–**d**) *In situ* SAXS analysis of thermal softening in hydrated rice culm cell walls. SAXS profiles of hydrated rice culm samples were collected at 25°C, 60°C, and 90°C. Representative background-subtracted equatorial *q*–*I* (**b**) and Kratky (**c**) plots of culm cell walls from WT1 and *OsC3′H1*-KD plants, alongside the relative scattering intensity increase (Δ*I*) compared with 25°C across all examined lines (**d**), are shown. Additional *q*–*I* and Kratky profiles across all lines are provided in **Figures S7** and **S8**. The Os4CL4-KO line was omitted from this analysis due to insufficient X-ray scattering contrast in its SAXS profiles, whereas WT2 was excluded because its SAXS profiles were virtually identical to those of WT1 (Figure 3). In **a** and **d**, data represent means ± standard deviation from three analytical runs (*n* = 3). Asterisks (*) indicate a statistically significant difference from the respective wild-type controls (Student’s *t*-test, \*\**p* < 0.01, \**p* < 0.05). WT1 and WT2, wild-type control lines; *OsCAld5H1*-KO, *OsCAld5H1*-knockout line; *OsCAld5H1*-OX, *OsCAld5H1*-overexpressing line; *OsC3′H1*-KD, *OsC3′H1*-knockdown line; *Os4CL3*-KO, *Os4CL3*-knockout line; *Os4CL4*-KO, *Os4CL4*-knockout line; *OsHCALDH2/3*-DKO, *OsHCALDH2-* and *OsHCALDH3*-double-knockout line. WT2 and *OsHCALDH2/3*-DKO, and WT1 and all other rice lines, were grown side-by-side.

### *In situ* SAXS profiling of thermal softening

To probe thermal softening of rice cell walls, fully hydrated intact culm samples with erased thermal history were placed in a water-filled liquid cell, and SAXS profiles were collected sequentially across elevated temperatures (25 °C, 60 °C, and 90 °C) (Horiyama et al., 2022). The *Os4CL4*-KO line was omitted from this analysis due to insufficient X-ray scattering contrast in its SAXS profiles (**Figure 3**), which precluded a reliable comparative evaluation against the remaining lines. Additionally, WT2 was excluded because its SAXS profiles were virtually identical to those of WT1.

The equatorial scattering intensity profiles (*q*–*I* plots) at the *q* < 0.2 Å⁻¹ increased in parallel with the rise in temperature across all rice lines (**Figure 4b; Figure S7**), which was further visualized by converting the *q*–*I* profiles into Kratky plots (**Figure 4c; Figure S8**). This qualitatively indicates that the electron density contrast between CMFs and lignin– hemicellulose matrix increases at elevated temperatures, mechanistically reflecting the thermal softening of matrix polymers, most likely lignin (Horiyama et al., 2022), as further discussed below. To compare the thermal softening behavior of lignin- and FA-altered rice cell walls, we calculated the intensity increase (Δ*I*) relative to 25 °C at the peak maximum of the Kratky plots (**Figure 4d**). Consequently, Δ*I* values were significantly elevated for *OsC3′H1*-KD at 60 °C, as well as all FA-depleted lines (*OsC3′H1*-KD, *Os4CL3*-KO and *OsHCALDH2/3*-DKO) at 90 °C compared to those of the WT1 control. In contrast, the the G-lignin-enriched *OsCAld5H1*-KO showed Δ*I* values comparable to those of WT1, whereas the S-lignin-enriched *OsCAld5H1*-OX line exhibited an apparent upward trend, albeit without statistical significance.

Overall, FA-depleted lines exhibited significantly enhanced enzymatic saccharification—with the notable exception of the *Os4CL4*-KO mutant—and accelerated thermal softening relative to G/S-lignin-altered lines. This implies that the disrupted lignocellulose assembly induced by reduced FA cross-linking can broadly translate into distinct biomass properties, although this effect can be critically modulated by other structural factors, as discussed below.

## Discussion

The role of FA in cross-linking AX and lignin in commelinid monocot cell walls, particularly in grasses, has been extensively investigated. This robust cross-linking within the cell walls is thought to be the cause of the limited digestibility of grass biomass in both ruminants and industrial biorefining (Buanafina, 2009; Ralph, 2010; de Oliveira et al., 2015; Hatfield et al., 2017; de Souza et al., 2018; Mnich et al., 2020; Chandrakanth et al., 2023; Yamamoto et al., 2024; Yang et al., 2024), and may explain why grasses require less lignin to support their cell wall structure than typical eudicot and gymnosperm species (Smith and Ralph, 2024). Nevertheless, the structural basis by which FA-mediated cross-linking dictates the overarching lignocellulose supramolecular assembly and macroscopic biomass properties has remained elusive. Our comparative ultrastructural analysis of lignin- and FA-altered rice lines particularly highlights the pivotal role of FA cross-linking in dictating grass cell wall architecture and biomass properties (**Figure 5**), solidifying its position as a prime structural target for advancing grass biomass utilization and crop design.

**Figure 5.**
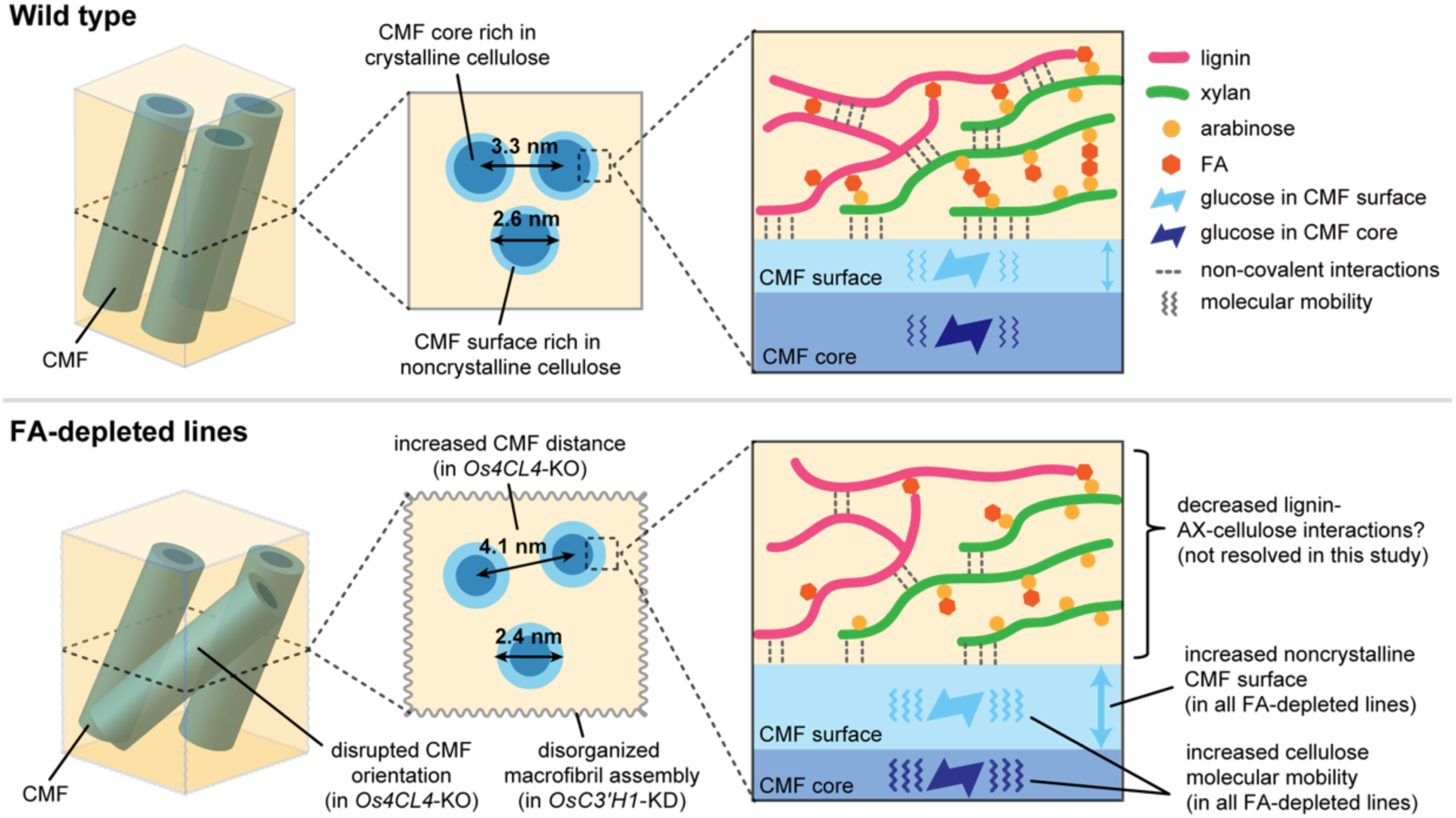
Proposed model of disrupted lignocellulose assembly in ferulate-depleted rice cell walls. The depletion of ferulate (FA)-mediated cross-linking within the lignin–arabinoxylan (AX) matrix drives a loosening of the supramolecular network. This loosening results in an increased proportion of noncrystalline cellulose microfibril (CMF) surfaces and elevated cellulose molecular mobility within the CMFs. These disruptions in cellulose crystal structure and mobility are likely triggered by diminished AX–AX and AX–lignin interactions, which may subsequently alter AX–cellulose and lignin–cellulose associations. Ultimately, this molecular- scale disorder can cascade into structural defects at the nano- and mesoscales, leading to disorganized CMF packing and orientation, and extending to broader disruptions at the macrofibrillar level.

Combined ssNMR and WAXD analyses indicate that depleting FA cross-linking drives supramolecular disorder of lignocellulose within the cell walls. Across all FA-reduced lines (*OsC3′H1*-KD, *Os4CL3*-KO, *Os4CL4*-KO, and *OsHCALDH2/3*-DKO), we observed consistently lower cellulose crystallinity and elevated cellulose molecular mobility, whereas no such changes were detected in rice lines with altered G/S lignin unit composition (*OsCAld5H1*-KO and *OsCAld5H1*-OX) (**Figure 2**). Furthermore, SAXS analysis detected disruptions in meso-scale CMF packing structure in not all but specific FA-depleted lines, i.e., *Os4CL4*-KO and *OsC3’H1*-KD (**Figure 3**), suggesting that molecular-scale lignocellulose perturbations induced by reduced FA cross-linking can occasionally cascade into nano- and meso-scale cell wall defects (**Figure 5**). Importantly, however, the notable reduction in total lignin content observed in our rice mutant and transgenic lines (**Table 1**) warrants careful consideration, as severe lignin depletion is known to alter cellulose assembly within plant cell walls (Ruel et al., 2009; Liu et al., 2016). In our study, however, close examination of the variations in lignin content—such as those between the two WT control lines (WT1 vs. WT2) or among the G/S-lignin- and the FA-altered lines with similar lignin contents (e.g., *OsCAld5H1*-OX vs. *Os4CL4*-KO and *OsHCALDH2/3*-DKO) (**Table 1**)—revealed no clear correlation with supramolecular structural parameters (**Figures 2, 3**). Nevertheless, potential synergistic interactions between lignin reduction and FA depletion cannot be entirely excluded.

Considering the current model of lignocellulose assembly wherein hemicelluloses bridge lignin and cellulose, and the latter two polymers retain limited direct contacts (Terashima et al., 2009; Kang et al., 2019; Terrett and Dupree, 2019; Kirui et al., 2022), disrupting FA-mediated AX–AX, AX–lignin, and lignin–lignin cross-linking likely cascades into perturbed AX–cellulose and cellulose–cellulose interactions, ultimately loosening cellulose crystalline structure and molecular constraints, occasionally extending to mesoscale CMF packing defects (**Figure 5**). Supporting this model, recent solid-state NMR analyses of *Brachypodium distachyon* stems revealed that feruloylated AX chains extensively interact with cellulose microfibrils, forcing lignin to bind to CMF associated with these AX chains (Duan et al., 2021). Gao et al. (2020) reported that AX–cellulose interfaces in the cell walls of the grass *Sorghum bicolor* are dominated by relatively weak interactions compared to the more robust xylan-cellulose interactions found in eudicot and softwood cell walls (Simmons et al., 2016; Terret et al., 2019). It is therefore plausible that FA-mediated AX–lignin cross-linking plays a compensatory role in fortifying the structural integrity of the grass cell walls.

Such disruptions in the lignocellulose supramolecular assembly of FA-depleted lines may broadly translate into distinct macroscopic biomass properties (**Figure 4**). Indeed, compared to the WT controls and G/S-lignin-altered mutants, most of the examined FA-depleted lines exhibited substantially higher saccharification efficiencies both with and without alkaline pretreatment. However, consistent with our earlier report (Affifi et al., 2022), the *Os4CL4*-KO line—which displayed one of the most severely disrupted lignocellulose assembly among the FA-depleted lines (**Figures 2, 3**)—did not show any improvement in saccharification efficiency without alkaline pretreatment, although it achieved the highest efficiency following pretreatment (**Figure 4**). This notable exception precludes the definitive conclusion that disrupted lignocellulose assembly in FA-depleted lines uniformly translates into improved saccharification. Our previous NMR study detected a substantial depletion of tricin and a concomitant increase in *p*-coumarate units within the *Os4CL4*-KO lignin (Affifi et al., 2022). Because such alterations in tricin and *p*-coumarate modifications can substantially affect the macromolecular structure and properties of grass lignin (Lam et al., 2024, 2026), it is plausible that these resulting modifications elevate lignin recalcitrance, offsetting the benefits of reduced FA cross-linking. Future investigations will be required to deconvolute these divergent structural effects.

All examined FA-depleted rice lines exhibited markedly enhanced thermal softening of lignocellulose (**Figure 4**), corroborating our notion that the disruption of FA-mediated cross- linking accelerates thermal dissociation of lignocellulose. Within the temperature range investigated here, this acceleration is most likely driven by lignin softening, given that the thermal softening temperature or glass transition temperature (*T*_g_) of hydrated lignin typically falls within this window (60–100 °C) (Olsson and Salmén, 1992; Furuta et al., 2008; Börcsök and Pásztory, 2021). Thus, disrupting FA-mediated cross-linking likely accelerates the thermal softening of lignin, either by directly lowering the *T*_g_ of lignin polymers, indirectly by reducing physical constraints imposed by FA-mediated cross-linking, or a combination of both. The extent to which lignin structure governs its thermomechanical properties remains debated (Börcsök and Pásztory, 2021). Comparisons between softwood (G) and hardwood (G/S) lignins suggest that a higher S/G ratio lowers lignin *T*_g_ (Olsson and Salmén, 1992; Furuta et al., 1997; 2008; Placet et al., 2007). In contrast, a study on transgenic poplar lines with varying G/S ratios detected no significant influence of the G/S ratio on wood thermal softening (Horvath et al., 2011). Here, the S-lignin-enriched *OsCAld5H1*-OX line exhibited a trend toward accelerated softening relative to WT controls and the G-lignin-enriched *OsCAld5H1*-KO line but, unlike the FA-depleted lines, this effect was not statistically significant (**Figure 4**). Future investigations must explicitly compare the thermal properties of isolated lignins with their corresponding intact cell walls to clarify the relationship between lignin structure and biomass thermal properties.

It is also noteworthy that the *OsC3′H1*-KD line, which exhibits a severe reduction in FA cross-linking, possesses altered lignin substantially enriched in H units (**Figure 1**) (Takeda et al., 2018; Takeda-Kimura et al., 2025). Previously, the *Arabidopsis med5a/5b ref8* mutant, which accumulates unusually high levels of H-lignin, was shown to exhibit improved enzymatic saccharification (Bonawitz et al., 2014; Shi et al., 2016). Furthermore, X-ray microdiffraction analysis suggested that this *Arabidopsis* mutant is susceptible to CMF disorganization upon exposure to moisture (Liu et al., 2016). Since the eudicot *Arabidopsis* naturally lacks FA-mediated cell wall cross-linking, these phenotypes likely arise from the extensively altered polymeric characteristics of lignin heavily enriched in H units compared with normal G/S lignin (Ralph et al., 2006; Ziebell et al., 2010; Bonawitz et al., 2014; Takeda et al., 2018). Thus, the disrupted lignocellulose molecular assembly and altered biomass properties observed in our *OsC3′H1*-KD line may stem from a synergistic effect between H lignin enrichment and FA depletion, a hypothesis that warrants further experimental exploration.

In contrast to the FA-depleted lines investigated in this study, or previously reported rice mutants with extensively altered lignin structures (e.g., those producing abnormal aldehyde and benzodioxane units) (Martin et al., 2019; 2023), the present G- or S-lignin enriched lines exhibited no major alterations in lignocellulose assembly, suggesting that the G/S ratio exerts minimal influence on cell wall architecture, at least within the G/S ratio range and structural parameters investigated in this study. Meanwhile, recent ssNMR studies using ^13^C-labeled cell wall samples have enabled the direct detection and quantitative assessment of non-covalent polysaccharide–polysaccharide (García et al., 2011; Dupree et al., 2015; Simmons et al., 2016; Terrett et al., 2019; Gao et al., 2020; Duan et al., 2021) and even more elusive lignin– polysaccharide interactions (Kang et al., 2019; Kirui et al., 2022; Addison et al., 2024; Hu et al., 2025, 2026; Xiao et al., 2025) within native cell walls. Notably, several studies have indicated that lignin associates with both hemicelluloses and cellulose most prominently via its methoxy groups (Kang et al., 2019; Kirui et al., 2022; Addison et al., 2024; Hu et al., 2025, 2026). In particular, Xiao et al. (2025) revealed that an increased S/G ratio (and the concomitant enrichment in methoxy groups) greatly stabilizes the lignin–polysaccharide interface in *Arabidopsis* inflorescence stems. Consequently, the application of these advanced NMR methods is required to resolve the specific polysaccharide–polysaccharide and lignin– polysaccharide interactions driving the FA- and lignin-induced alterations in the cell wall architecture observed in our rice lines. Such studies will advance our understanding of the still-elusive supramolecular structure of plant cell walls and its contribution to their function and utility.

### Experimental procedures

#### Plant materials

Previously characterized rice (*Oryza sativa* L. cv. Nipponbare) lines were utilized in this study: *OsCAld5H1*-KO (identical to *OsCAld5H1*-KO-a-9-2; Takeda et al., 2019a), *OsCAld5H1*-OX (*OsCAld5H1*-OX-a; Takeda et al., 2017), *OsC3′H1*-KD (*OsC3′H1*-KD-a; Takeda et al., 2018), *Os4CL3*-KO and *Os4CL4*-KO (*os4cl3-a* and *os4cl4-a*; Afifi et al., 2022), and *OsHCALDH2/3*- DKO (*oshcaldh2-2 oshcaldh3-3*; Yamamoto et al., 2024). All lines, along with their corresponding wild-type controls (WT1 and WT2), were grown to maturity under greenhouse conditions (**Table S1**) (Lam et al., 2017). Mature aerial tissues were harvested and dried at 45 °C for 3 days. To prepare extractive-free CWR for chemical analyses, WAXD, ssNMR, and saccharification assays, dried culms were cut into ∼5-mm segments and homogeneously pulverized using a TissueLyser (Qiagen, Hilden, Germany) operated at 25 Hz for 3 min per 600-mg sample. The pulverized tissue was subjected to sequential solvent extraction as described previously (Yamamura et al., 2012). For SAXS measurements, intact culm segments (1.0 cm in length) were excised from the main tillers within a 2.0–3.0 cm region above the second internode (toward the first internode). Each cylindrical segment was longitudinally split open and trimmed into a 1.0 × 1.0 cm square prior to further processing, as detailed below.

#### Chemical analyses

The Klason lignin assay (Hatfield et al., 1994), analytical thioacidolysis (Yamamura et al., 2012), quantification of cell wall-bound FA and *p*-coumarate via alkaline hydrolysis (Yamamura et al., 2011), quantification of arabinoxylan-bound FA via methanolic acidolysis (Lam et al., 2024) using synthetic **MeAra-FA** (Hatfield et al., 1991; Helm et al., 1992), and neutral sugar analysis (Lam et al., 2017) were all performed as previously described.

#### WAXD

WAXD analysis of CWR samples was performed using a Rigaku Ultima-IV diffractometer (Rigaku, Tokyo, Japan) operating with nickel-filtered CuKα radiation (λ = 1.54 Å) at 40 kV and 40 mA. Samples were mounted on a copper sample holder, and diffraction intensity profiles were recorded at 20 °C over a 2*θ* range of 5° to 30° with a step size of 0.1° and a dwell time of 15 s per step. Multi-peak deconvolution of the WAXD profiles was conducted in Igor Pro 9 (WaveMetrics, Portland, OR, USA) by fitting Voigt functions to the 200 reflection, the combined 110/1 1 0 reflections, and the amorphous background contribution. The cell wall crystallinity (*CrI*) was calculated from the maximum intensity of the 200 reflection (*I*_200_, 2*θ* ≈ 22°) and the minimum intensity between the 200 and 110/1 1 0 peaks (*I*_am_, 2θ ≈ 18°) according to the following equation (Segal et al., 1959):

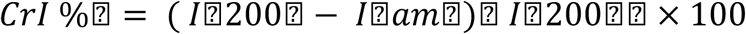

The apparent cellulose crystal size (D) was determined from the FWHM of the 200 peak (2*θ* = ca. 22°) using the following equation (Scherrer, 1918):

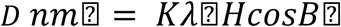

where *K* is constant (0.9), *λ* is wavelength (CuKα = 0.1542 nm), H is the FWHM of the 200 peak (rad.), and *B* is the peak position of the 200 peak.

#### ssNMR

ssNMR analysis of rice CWRs was performed on a Bruker Biospin Avance III 800US system (Bruker Biospin, Billerica, MA, USA) with a 4-mm double resonance MAS probe and a ZrO_2_ rotor with a KelF-made cap. The ^13^C CP MAS spectra were collected at 300K with 12k Hz MAS frequency, 2.5 ms contact time, and 4.0 s recycling delay of 4.0 s to preferentially detect rigid cellulose (Martin et al., 2019; 2023). ^13^C chemical shifts were externally referenced to CH_3_ signal of silicon rubber at 1.4 ppm on the tetramethylsilane scale. The *T*_1_ measurements were conducted using the Torchia pulse sequence (Torchia, 1978) with a CP contact time of 2.5 ms and 11 or 12 τ delay times between 0.01 and 64 s (Martin et al., 2019; 2023). The obtained τ-dependent signal decay *I*(τ) was fitted using Igor Pro 9 software (WaveMetrics, Inc., Portland, OR, USA) with the following double exponential equation:

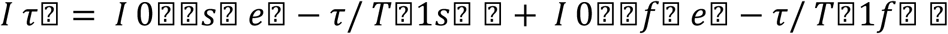

where *T*_1s_ and *T*_1f_ are two independent *T*_1_ for slower and faster relaxing components (*T*_1s_ > *T*_1f_), and *I* (0)_s_ and *I* (0)_f_ are their weighing fractions (Martin et al., 2019; 2023).

#### SAXS measurements

SAXS measurements on rice culms were performed at the SPring-8 synchrotron radiation facility (RIKEN, Hyogo, Japan) using X-rays with a wavelength (λ) of 1.0 Å and a Pilatus hybrid pixel array detector (Dectris Inc., Switzerland). Incident and transmitted X-ray fluxes were monitored using upstream and downstream ionization chambers, respectively, to correct the scattering data for sample absorption. The magnitude of the scattering vector, *q*, is defined as *q* = 4πsin(*θ*)/λ, where 2*θ* is the scattering angle. For dry-state rice culms, SAXS measurements were conducted on the BL40B2 beamline. Samples were placed in quartz cells with a 2-mm path length, oriented such that both the lumen plane and fiber direction were perpendicular to the X-ray beam. Data were collected at a sample-to-detector distance of 1.2 m, covering a *q*-range of 0.01–0.7 Å⁻¹ with an exposure time of 10 s. For *in situ* measurements of hydrated rice culms, SAXS experiments were carried out on the BL19B2 beamline equipped with an mK2000 temperature controller (Instec Inc., USA). Prior to analysis, air-dried culm sections (1 × 1 cm) were vacuum-infiltrated with distilled water to complete saturation, heat-treated at 100 °C for 10 min to erase thermal history, and subsequently cooled to room temperature. Samples were mounted in an in-house liquid cell and heated stepwise to 25, 60, and 90 °C, equilibrating for 4 min at each target temperature prior to data collection. Data were acquired at a sample-to-detector distance of 1.7 m, covering a *_q_*-range of 0.01–0.5 Å⁻¹ with an exposure time of 60 s.

#### SAXS data analysis

SAXS data reduction and analysis were performed using the *pyFAI* (Ashiotis et al., 2015) and FabIO (Knudsen et al., 2013) Python packages, following previously reported procedures (Penttilä et al., 2019; Horiyama et al., 2022) with minor modifications. Briefly, the obtained 2D scattering patterns were corrected by its absorbance and background-subtracted, and *χ–I* plots were generated by integrating over a *q*-range of 0.075–0.25 Å⁻¹. From these profiles, two characteristic azimuthal angles were identified: the angle of maximum intensity (equatorial direction, defined as *χ* = 0°) and the angle of minimum intensity (isotropic background originating from the homogeneous cell wall matrix). For CMF orientation analysis, the obtained *χ–I* plots were subjected to multi-peak fitting using three Gaussian peaks to resolve one central radial peak and two symmetrical tangential peaks. The FWHM of the central radial peak was subsequently used to evaluate the overall orientational heterogeneity of CMFs (Lichtenegger et al., 1999). In parallel, *q–I* plots were generated by performing sector integration around these two angles using a 15° azimuthal window. The isotropic background profile was then subtracted from the equatorial 1D scattering data, and Kratky plots were constructed and smoothed using a Gaussian filter. For *in situ* SAXS measurements of hydrated rice culms, temperature-dependent structural alterations were evaluated by calculating the relative change in scattering intensity (Δ*I*) at the Kratky peak maximum at each temperature (25 °C, 60 °C, and 90 °C) with respect to the baseline intensity at 25°C. For dry-state culm samples, *q–I* scattering profiles were analyzed in SasView (Doucet et al., 2020) using the WoodSAS model equation (Penttilä et al., 2019):

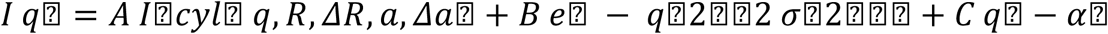

where *A*, *B*, *σ*, *C*, and *ɑ* are scaling and decay constants, *I*_cyl_ (*q*) represents the scattering intensity from an array of infinitely long cylinders corresponding to CMFs in the secondary cell wall (Penttilä et al., 2019). Here, *R* is the mean microfibril radius with a standard deviation of Δ*R*, and *a* is the center-to-center interfibrillar packing distance with a paracrystalline lattice distortion of Δ*_a_*. The Gaussian function centered at *q* = 0 Å^−1^ (associated with constant *B*) accounts for form-factor scattering from nanoscale features larger than individual CMFs. The power-law scattering term (associated with constant *C*) at low *q*, with α ≈ 4, is attributed to scattering from the surfaces of larger pores and cell lumina (Penttilä et al., 2019; 2020).

#### Enzymatic saccharification assay

Enzymatic saccharification assays using a cellulolytic enzyme cocktail both with and without mild alkaline pretreatment (312.5 mM NaOH, 90 °C, 3 h) were performed as previously described (Takeda et al., 2019b; Yamamoto et al., 2024).

## Accession numbers

Rice genes investigated in this article can be found under accession numbers: LOC_Os10g36848 (*OsCAld5H1*; Takeda et al., 2017), LOC_Os05g41440 (*OsC3′H1*; Takeda et al., 2018), LOC_Os02g08100 (*Os4CL3*; Afifi et al., 2022), LOC_Os06g44620 (*Os4CL4*; Afifi et al., 2022), LOC_Os01g40870 (*OsHCALDH2*; Yamamoto et al., 2024) and LOC_Os06g39230 (*OsHCALDH3*; Yamamoto et al., 2024).

## Acknowledgments

We thank Prof. John Ralph and Dr. Steven D. Karlen (University of Wisconsin–Madison), and Dr. Richard Helm (Virginia Tech) for providing the synthetic **MeAra-FA** standard and guidance on mild acidolysis, as well as Prof. Hironori Kaji and Ms. Ayaka Maeno (Kyoto University) for assistance with NMR analyses. We also thank Novozymes Japan Ltd. (Chiba, Japan) for providing the enzyme cocktails used in the saccharification assays. This work was supported in part by JSPS KAKENHI (Grant Nos. JP20H03044 and JP24K01827). This study was partially conducted using the greenhouse and GC-/LC-MS facilities of the Development and Assessment of Sustainable Humanosphere/Forest Biomass Analytical System (DASH/FBAS, Research Institute for Sustainable Humanosphere, Kyoto University); the NMR spectrometer at the International Joint Usage/Research Center (iJURC, Institute for Chemical Research, Kyoto University); and SAXS beamlines BL40B2 and BL19B2 at SPring-8 (RIKEN), with the approval of the Japan Synchrotron Radiation Research Institute (JASRI; Proposal Nos. 2020A1592, 2021A1384, 2021B1682, 2022B1462, 2023A1480, and 2024A1728).

## Author contributions

S.Y., T.I., and Y.T. conceived the study and wrote the manuscript with input from all authors. S.Y., R.K., Ka.K., Ke.K., T.I., T.U., and Y.T. designed the experiments. S.Y., O.A.A., P.J., R.K., Ka.K., Ke.K., T.I., and Y.T. performed the experiments and analyzed the data.

## Data Availability Statement

The authors confirm that the data supporting the findings of this study are available within the article and its supplementary materials.

## Supporting Information

Additional supporting information can be found online in the Supporting Information section.

**Figure S1. WAXD profiles of rice cell walls.**

**Figure S2. ssNMR spectra of rice cell walls.**

**Figure S3. SAXS profiles of dry-state rice cell walls.**

**Figure S4. *χ-I* plots of dry-state rice cell walls.**

**Figure S5. *q-I* plots of dry-state rice cell walls.**

**Figure S6. Kratky plots of dry-state rice cell walls.**

**Figure S7. *q-I* plots of hydrated rice cell walls.**

**Figure S8. Kratky plots of hydrated rice cell walls.**

**Table S1. Growth characteristics of lignin- and FA-altered rice.**

**Table S2. Signal assignments for ssNMR spectra of rice cell walls.**

**Table S3. ssNMR-derived relaxation data of rice cell walls.**

**Table S4. WoodSAS fitting data of drya-state rice cell walls.**

