## Supporting Information for "Ultrastructural analysis of engineered rice lines reveals ferulate cross-linking as a key factor mediating lignocellulose supramolecular assembly in grass cell walls"

### List of Materials

| | |
|---|---|
| **Figure S1.** | **WAXD profiles of rice cell walls.** |
| **Figure S2.** | **ssNMR spectra of rice cell walls.** |
| **Figure S3.** | **SAXS profiles of dry-state rice cell walls.** |
| **Figure S4.** | ***χ*-*I* plots of dry-state rice cell walls.** |
| **Figure S5.** | ***q*-*I* plots of dry-state rice cell walls.** |
| **Figure S6.** | **Kratky plots of dry-state rice cell walls.** |
| **Figure S7.** | ***q*-*I* plots of hydrated rice cell walls.** |
| **Figure S8.** | **Kratky plots of hydrated rice cell walls.** |
| **Table S1.** | **Growth characteristics of lignin- and FA-altered rice.** |
| **Table S2.** | **Signal assignments for ssNMR spectra of rice cell walls.** |
| **Table S3.** | **ssNMR-derived relaxation data of rice cell walls.** |
| **Table S4.** | **WoodSAS fitting data of drya-state rice cell walls.** |

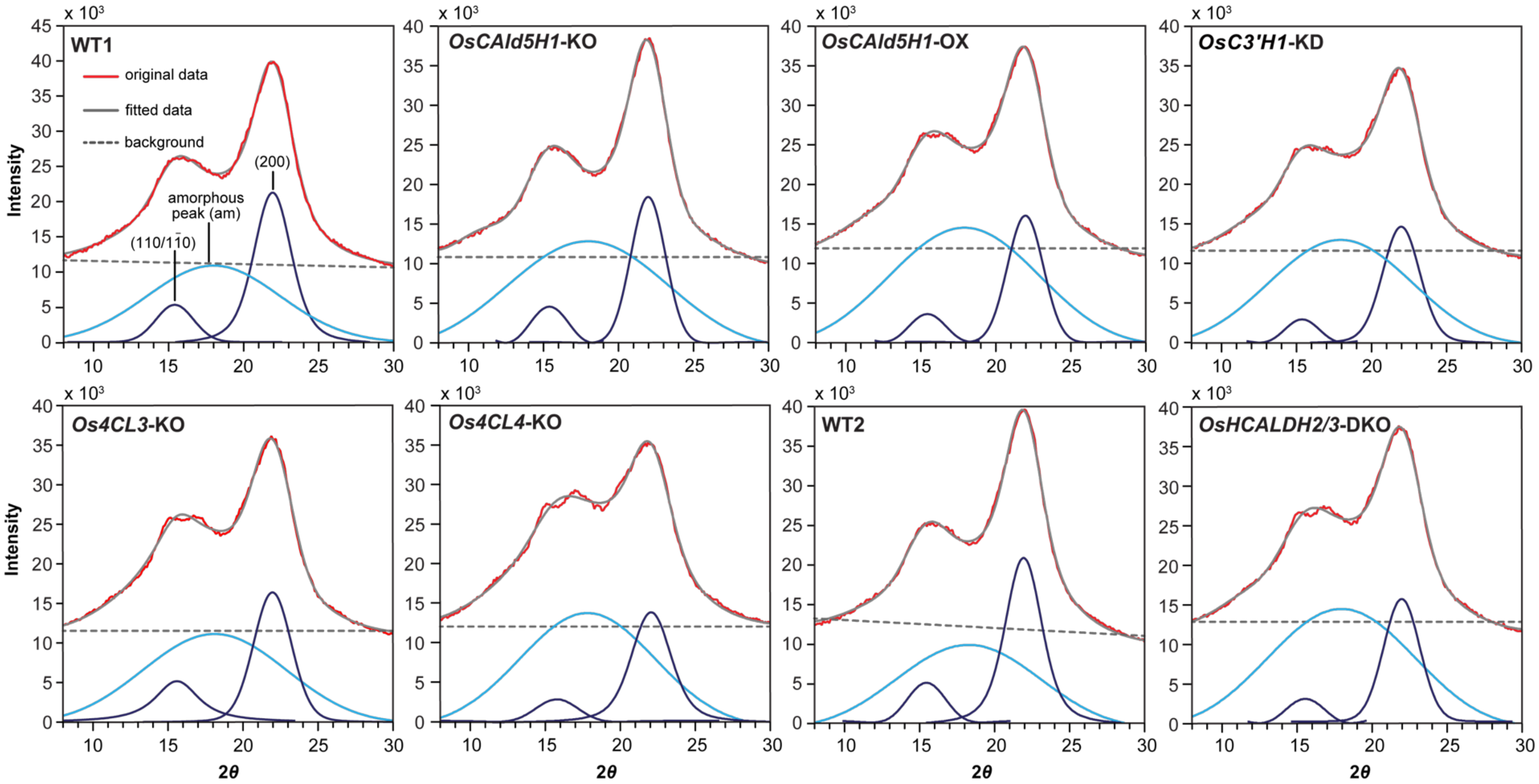


**Figure S1. WAXD profiles of rice cell walls.**

Representative wide-angle X-ray diffraction (WAXD) profiles of culm cell walls are shown. The experimental WAXD profiles (red line) were deconvoluted via multi-peak fitting (gray line) into an amorphous contribution (light blue line) and crystalline cellulose reflections corresponding to the (200) and $(110/1\bar{1}0)$ lattice planes (dark blue lines). The dashed gray line indicates the baseline subtracted prior to fitting. WT1 and WT2, wild-type control lines; *OsCAld5H1*-KO, *OsCAld5H1*-knockout line; *OsCAld5H1*-OX, *OsCAld5H1*-overexpressing line; *OsC3'H1*-KD, *OsC3'H1*-knockdown line; *Os4CL3*-KO, *Os4CL3*-knockout line; *Os4CL4*-KO, *Os4CL4*-knockout line; *OsHCALDH2/3*-DKO, *OsHCALDH2*- and *OsHCALDH3*-double-knockout line. The WT2 and *OsHCALDH2/3*-DKO lines, as well as WT1 and all other transgenic lines, were grown side-by-side.

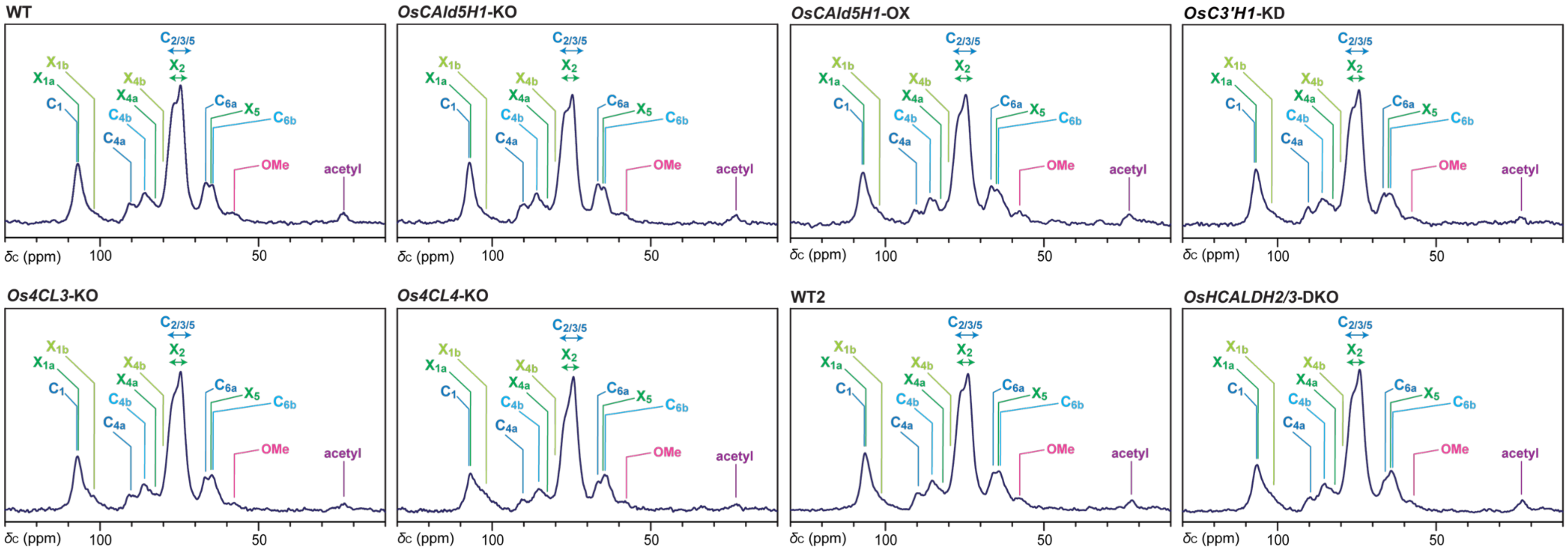


**Figure S2. ssNMR spectra of rice cell walls.**

[13]C magic-angle-spinning (MAS) NMR spectra were collected using [1]H–[13]C cross-polarization (CP). Peak assignments are provided in **Table S2**. WT1 and WT2, wild-type control lines; *OsCAld5H1*-KO, *OsCAld5H1*-knockout line; *OsCAld5H1*-OX, *OsCAld5H1*-overexpressing line; *OsC3'H1*-KD, *OsC3'H1*-knockdown line; *Os4CL3*-KO, *Os4CL3*-knockout line; *Os4CL4*-KO, *Os4CL4*-knockout line; *OsHCALDH2/3*-DKO, *OsHCALDH2*- and *OsHCALDH3*-double-knockout line. The WT2 and *OsHCALDH2/3*-DKO lines, as well as WT1 and all other transgenic lines, were grown side-by-side.

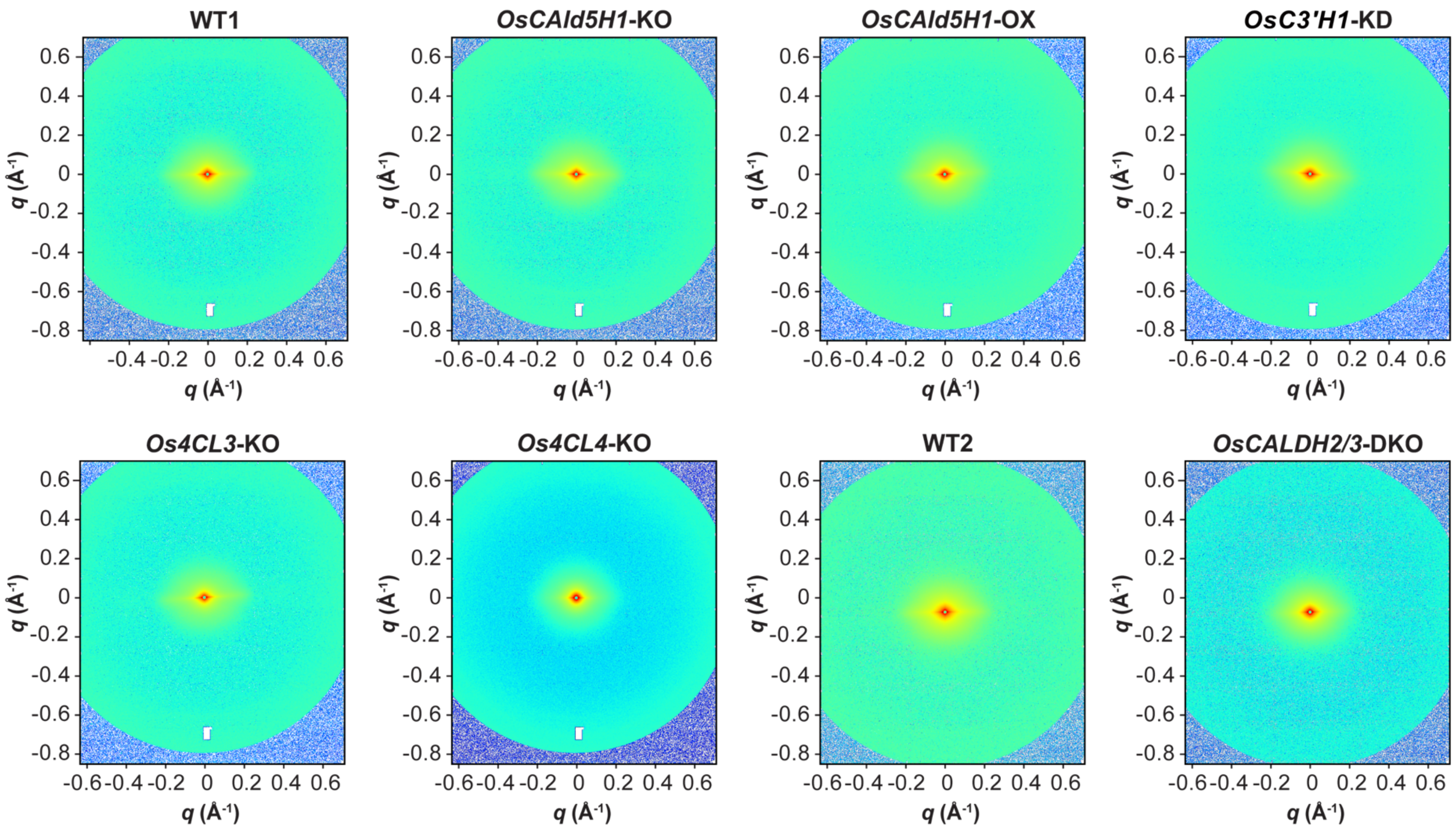


**Figure S3. 2D SAXS profiles of dry-state rice cell walls.**

Two-dimensional (2D) SAXS patterns were recorded from rice culm samples positioned such that the incident X-ray beam was directed perpendicular to both the longitudinal fiber axis and the lumen surface. WT1 and WT2, wild-type control lines; *OsCAld5H1*-KO, *OsCAld5H1*-knockout line; *OsCAld5H1*-OX, *OsCAld5H1*-overexpressing line; *OsC3'H1*-KD, *OsC3'H1*-knockdown line; *Os4CL3*-KO, *Os4CL3*-knockout line; *Os4CL4*-KO, *Os4CL4*-knockout line; *OsHCALDH2/3*-DKO, *OsHCALDH2*- and *OsHCALDH3*-double-knockout line. The WT2 and *OsHCALDH2/3*-DKO lines, as well as WT1 and all other transgenic lines, were grown side-by-side.

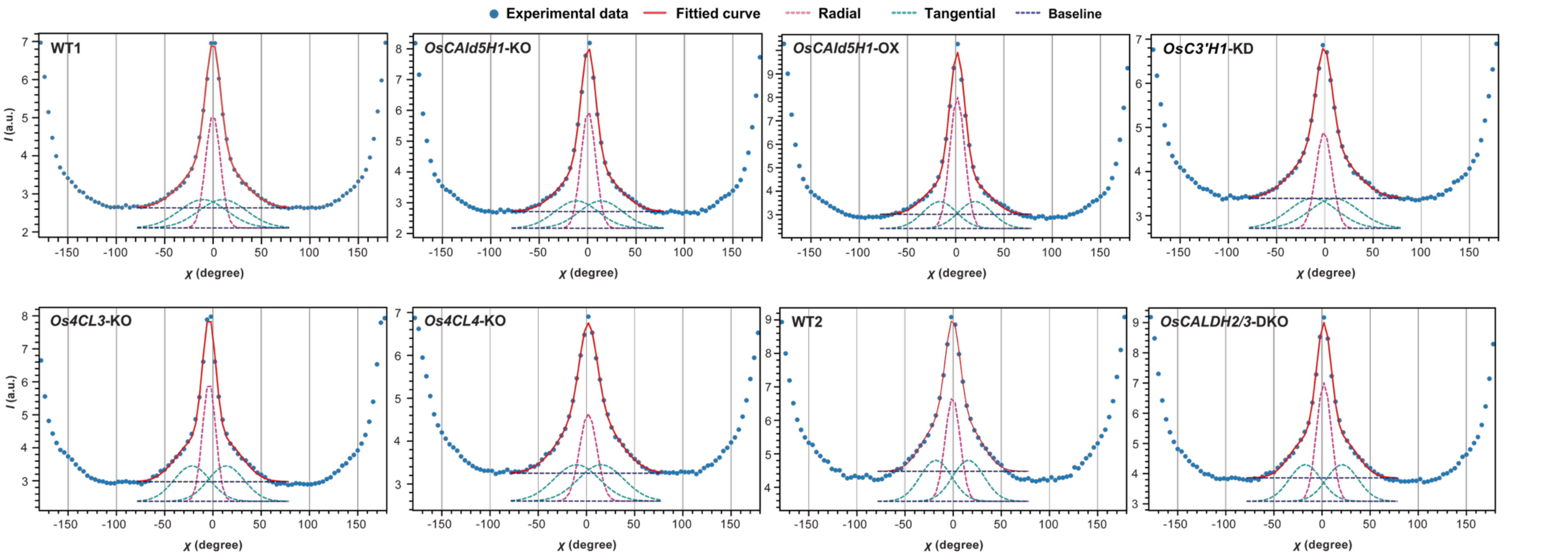


**Figure S4. *χ-I* plots of dry-state rice cell walls.**

The azimuthal intensity (*χ-I*) plot was fitted with three Gaussian functions across the range $q$ = 0.075-0.25 $Å^{-1}$ to resolve one radial and two tangential orientations. WT1 and WT2, wild-type control lines; *OsCAld5H1*-KO, *OsCAld5H1*-knockout line; *OsCAld5H1*-OX, *OsCAld5H1*-overexpressing line; *OsC3'H1*-KD, *OsC3'H1*-knockdown line; *Os4CL3*-KO, *Os4CL3*-knockout line; *Os4CL4*-KO, *Os4CL4*-knockout line; *OsHCALDH2/3*-DKO, *OsHCALDH2*- and *OsHCALDH3*-double-knockout line. The WT2 and *OsHCALDH2/3*-DKO lines, as well as WT1 and all other transgenic lines, were grown side-by-side.

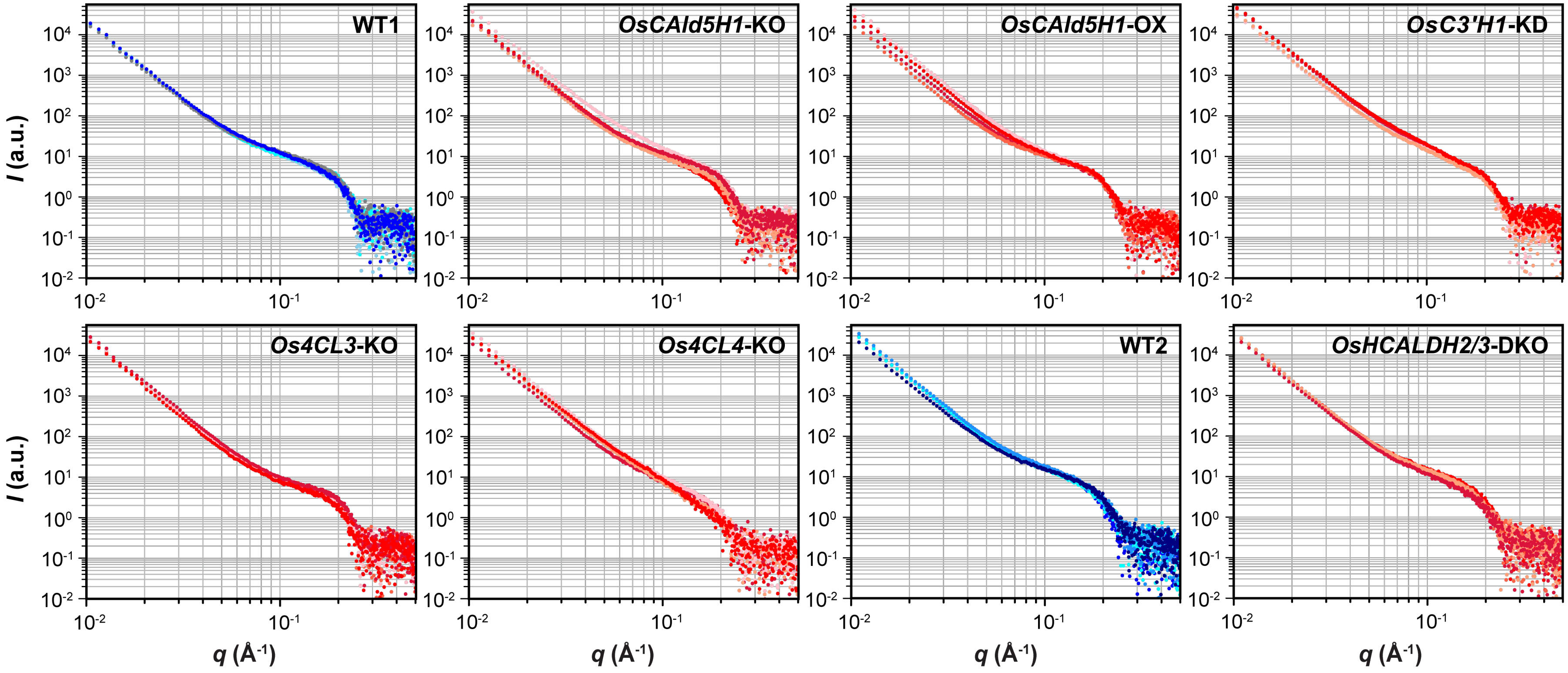


**Figure S5. *q-I* plots of dry-state rice cell walls.**

Background-subtracted equatorial *q-I* plots from the culm cell walls are shown. WT1 and WT2, wild-type control lines; *OsCAld5H1*-KO, *OsCAld5H1*-knockout line; *OsCAld5H1*-OX, *OsCAld5H1*-overexpressing line; *OsC3'H1*-KD, *OsC3'H1*-knockdown line; *Os4CL3*-KO, *Os4CL3*-knockout line; *Os4CL4*-KO, *Os4CL4*-knockout line; *OsHCALDH2/3*-DKO, *OsHCALDH2*- and *OsHCALDH3*-double-knockout line. The WT2 and *OsHCALDH2/3*-DKO lines, as well as WT1 and all other transgenic lines, were grown side-by-side.

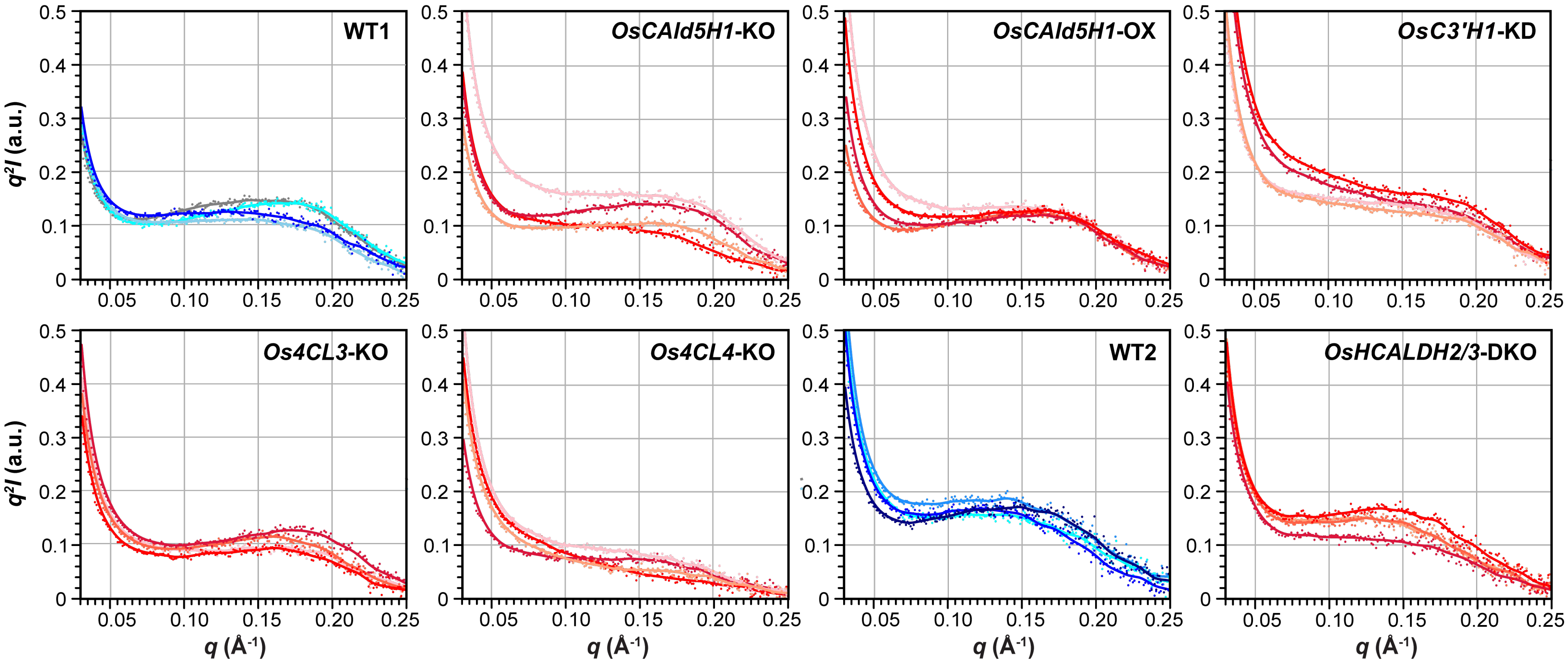


**Figure S6. Kratky plots of dry-state rice cell walls.**

The Kratky plots were constructed from the equatorial *q*-*I* plots from the culm cell walls. WT1 and WT2, wild-type control lines; *OsCAld5H1*-KO, *OsCAld5H1*-knockout line; *OsCAld5H1*-OX, *OsCAld5H1*-overexpressing line; *OsC3'H1*-KD, *OsC3'H1*-knockdown line; *Os4CL3*-KO, *Os4CL3*-knockout line; *Os4CL4*-KO, *Os4CL4*-knockout line; *OsHCALDH2/3*-DKO, *OsHCALDH2*- and *OsHCALDH3*-double-knockout line. The WT2 and *OsHCALDH2/3*-DKO lines, as well as WT1 and all other transgenic lines, were grown side-by-side.

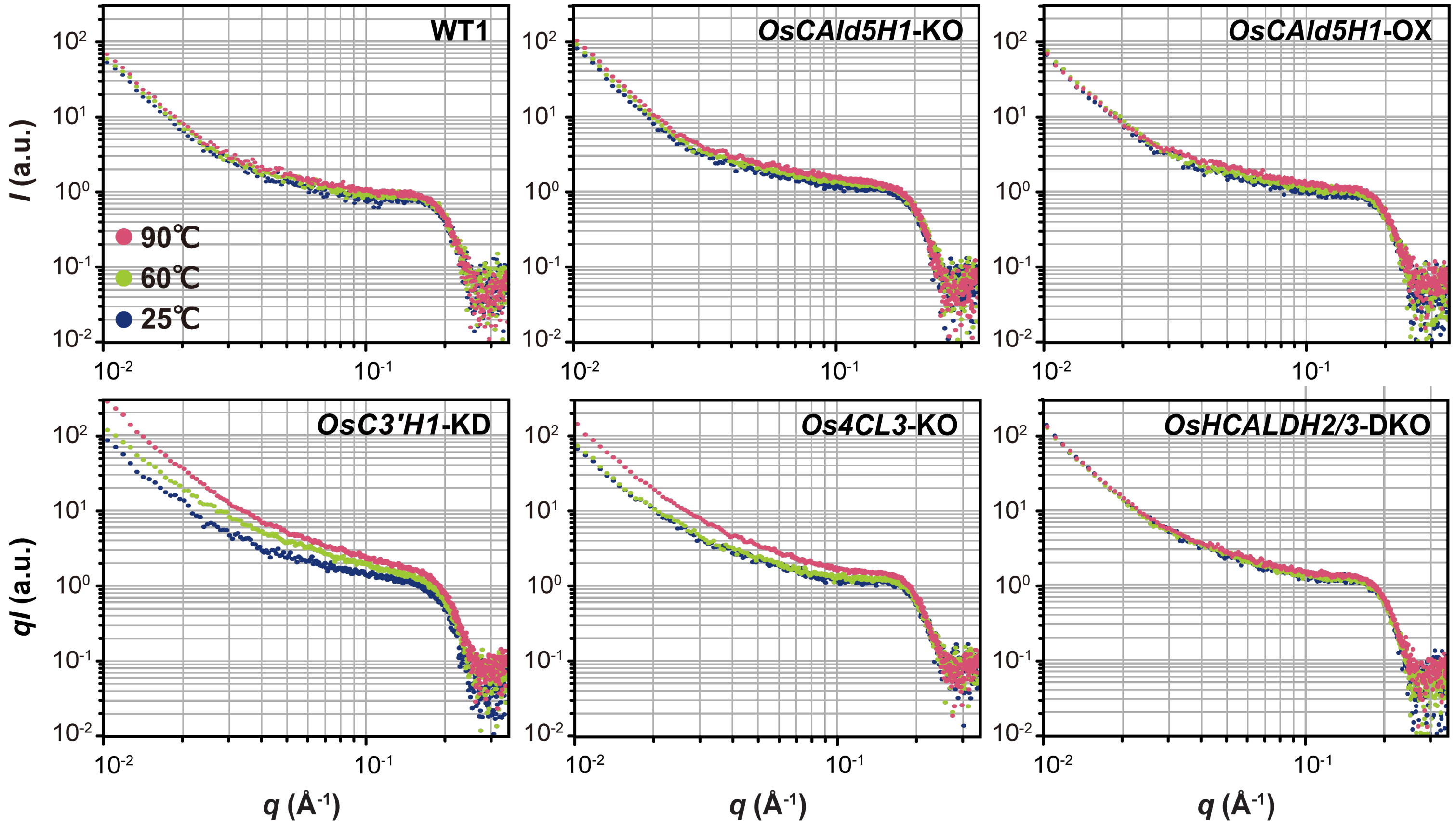


**Figure S7. *q-I* plots of hydrated rice cell walls.**

The Kratky plots were constructed from the equatorial *q-I* plots from hydrated culm cell walls at 25°C, 60°C, and 90°C. WT1, wild-type control lines; *OsCAld5H1*-KO, *OsCAld5H1*-knockout line; *OsCAld5H1*-OX, *OsCAld5H1*-overexpressing line; *OsC3'H1*-KD, *OsC3'H1*-knockdown line; *Os4CL3*-KO, *Os4CL3*-knockout line; *OsHCALDH2/3*-DKO, *OsHCALDH2*- and *OsHCALDH3*-double-knockout line.

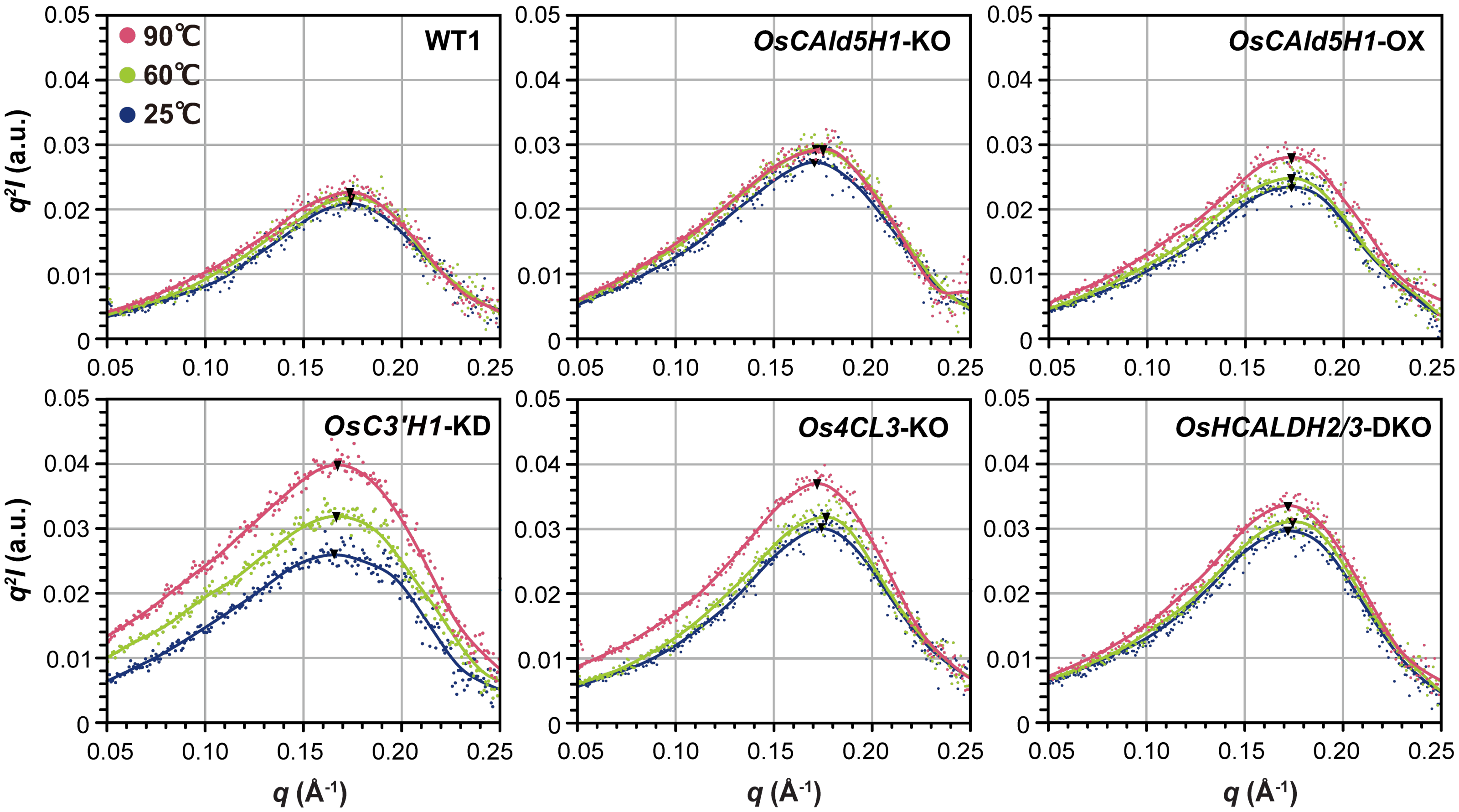


**Figure S8. Kratky plots of hydrated rice cell walls.**

The Kratky plots were constructed from the equatorial *q*-*I* plots from hydrated culm cell walls at 25°C, 60°C, and 90°C. WT1, wild-type control lines; *OsCAld5H1*-KO, *OsCAld5H1*-knockout line; *OsCAld5H1*-OX, *OsCAld5H1*-overexpressing line; *OsC3'H1*-KD, *OsC3'H1*-knockdown line; *Os4CL3*-KO, *Os4CL3*-knockout line; *OsHCALDH2/3*-DKO, *OsHCALDH2*- and *OsHCALDH3*-double-knockout line.

**Table S1. Growth characteristics of lignin- and FA-altered rice.**

| Trait | WT1 | *OsCAld5H1*-KO | *OsCAld5H1*-OX | *OsC3'H*-KD | *Os4CL3*-KO | *Os4CL4*-KO | WT2 | *OsHCALDH2/3*-DKO |
|---|---|---|---|---|---|---|---|---|
| Plant height (cm)[1] | 118.1 ± 5.5 | 114.8 ± 4.1 | **108.3 ± 6.0*** | 112.8 ± 5.1 | **109.6 ± 6.3*** | **90.3 ± 5.1**** | 116.3 ± 2.9 | **109.7 ± 5.6*** |
| Culm length (cm)[2] | 77.0 ± 6.2 | 74.6 ± 8.9 | **64.8 ± 4.1**** | 70.6 ± 4.2 | 76.0 ± 6.9 | **43.8 ± 7.4**** | 77.6 ± 2.3 | **73.8 ± 3.1*** |
| Panicle length (cm) | 24.0 ± 1.5 | **21.2 ± 1.7*** | 22.3 ± 1.6 | 22.7 ± 1.4 | 24.4 ± 5.6 | **16.8 ± 1.7**** | 17.4 ± 1.2 | **13.7 ± 2.9*** |
| Tiller number | 10.2 ± 1.5 | 10.5 ± 1.9 | **13.3 ± 1.9**** | 10.3 ± 2.2 | **13.8 ± 1.9**** | 12.2 ± 1.9 | 6.5 ± 1.6 | **7.8 ± 1.5*** |
| Biomass (g)[3] | 11.6 ± 0.7 | 12.8 ± 1.0 | 18.9 ± 5.3 | 10.3 ± 3.0 | **16.0 ± 0.3**** | 10.8 ± 1.3 | 15.8 ± 1.1 | 15.8 ± 1.6 |

[1]Length from the base of aerial part to the tip of top leaf. [2]Length from the base of aerial part to the base of panicle. [3]Dry weight of aerial parts excluding panicles. The values are means ± standard deviation of biological replicates ($n > 3$). Asterisks (*) and numbers in bold indicate significant difference from wild-type plants (Student's *t*-test, ***p* < 0.01, * *p* < 0.05). WT1 and WT2, wild-type control lines; *OsCAld5H1*-KO, *OsCAld5H1*-knockout line; *OsCAld5H1*-OX, *OsCAld5H1*-overexpressing line; *OsC3'H1*-KD, *OsC3'H1*-knockdown line; *Os4CL3*-KO, *Os4CL3*-knockout line; *Os4CL4*-KO, *Os4CL4*-knockout line; *OsHCALDH2/3*-DKO, *OsHCALDH2*- and *OsHCALDH3*-double-knockout line. The WT2 and *OsHCALDH2/3*-DKO lines, as well as WT1 and all other transgenic lines, were grown side-by-side.

**Table S2. Signal assignments for ssNMR spectra of rice cell walls.**

| Labels | Resonance (ppm) | Assignment |
|---|---|---|
| **$C_1$** | 106 | cellulose C1 |
| **$X_{1a}$** | $106^a$ | rigid/two-fold xylan C1 |
| **$X_{1b}$** | $103^a$ | mobile/three-fold xylan C1 |
| **$C_{4a}$** | 89 | crystalline/internal cellulose C4 |
| **$C_{4b}$** | 85 | noncrystalline/surface cellulose C4 |
| **$X_{4a}$** | $82^a$ | rigid/two-fold xylan C4 |
| **$X_{4b}$** | $77^a$ | mobile/three-fold xylan C4 |
| **$X_{2,3}$** | $75\text{-}72^a$ | xylan C2 and C3 |
| **$C_{2,3,5}$** | 75-72 | cellulose C2, C3 and C5 |
| **$C_{6a}$** | 66 | crystalline/internal cellulose C6 |
| **$X_5$** | $64^a$ | xylan C5 |
| **$C_{6a}$** | 63 | noncrystalline/surface cellulose C6 |
| **OMe** | 57 | lignin methoxy $CH_3$ |

[a]Not well resolved in this study. Chemical shift values are according to liretaures (Simmons et al. 2016; Martin et al. 2019; 2023).

**Table S3. CP $^{13}C$ spin-lattice relaxation data of rice cell walls.**

| | $C_1$ (106 ppm) | | $C_{4a}$ (89 ppm) | | $C_{4b}$ (85 ppm) | | $C_{6a}$ (66 ppm) | | $C_{6b}$ (63 ppm) | | OMe (57 ppm) | |
|---|---|---|---|---|---|---|---|---|---|---|---|---|
| $T_1$ (s) | *slow* | *fast* | *slow* | *fast* | *slow* | *fast* | *slow* | *fast* | *slow* | *fast* | *slow* | *fast* |
| **WT1** | 75.4 ± 3.9 | 0.06 ± 0.04 | 93.6 ± 1.7 | 0.22 ± 0.10 | 56.7 ± 3.6 | 0.05 ± 0.05 | 54.0 ± 7.2 | 0.12 ± 0.10 | 39.0 ± 6.0 | 0.69 ± 0.22 | 38.1 ± 5.6 | 1.78 ± 0.52 |
| ***OsCAld5H*-KO** | 71.5 ± 5.4 | 0.62 ± 0.43 | 91.1 ± 4.7 | 0.90 ± 0.29 | 51.4 ± 2.2 | 0.06 ± 0.03 | 54.3 ± 3.6 | 0.32 ± 0.14 | 46.2 ± 7.4 | 0.05 ± 0.03 | 43.6 ± 11.3 | 2.16 ± 0.62 |
| ***OsCAld5H1*-OX** | 79.4 ± 2.2 | 1.85 ± 0.48 | 86.3 ± 3.4 | 1.13 ± 0.35 | 53.4 ± 3.8 | 0.17 ± 0.15 | 50.0 ± 2.6 | 0.37 ± 0.09 | 37.7 ± 3.3 | 0.59 ± 0.10 | 41.3 ± 4.7 | 0.84 ± 0.24 |
| ***OsC3'H1*-KD** | 63.5 ± 2.9 | 0.08 ± 0.03 | 59.1 ± 3.8 | 1.02 ± 0.53 | 47.4 ± 4.3 | 0.03 ± 0.02 | 47.0 ± 6.6 | 0.13 ± 0.09 | 28.8 ± 4.3 | 0.23 ± 0.16 | 42.1 ± 7.6 | 0.17 ± 0.15 |
| ***Os4CL3*-KO** | 71.9 ± 3.4 | 0.05 ± 0.01 | 67.8 ± 4.7 | 0.06 ± 0.08 | 52.5 ± 1.0 | 1.06 ± 0.26 | 52.5 ± 4.8 | 0.12 ± 0.12 | 31.0 ± 3.8 | 0.78 ± 0.31 | 38.9 ± 5.4 | 2.24 ± 0.73 |
| ***Os4CL4*-KO** | 57.6 ± 8.1 | 0.09 ± 0.06 | 63.0 ± 10.0 | 0.07 ± 0.04 | 47.9 ± 3.9 | 0.05 ± 0.02 | 42.8 ± 8.5 | 0.13 ± 0.14 | 34.4 ± 6.0 | 0.95 ± 0.23 | 37.1 ± 9.4 | 1.07 ± 0.44 |
| **WT2** | 86.1 ± 3.0 | 0.03 ± 0.03 | 86.1 ± 1.8 | 0.11 ± 0.05 | 62.9 ± 1.9 | 0.28 ± 0.21 | 60.3 ± 6.2 | 1.70 ± 0.64 | 46.2 ± 6.7 | 1.53 ± 0.51 | 40.9 ± 8.3 | 2.77 ± 0.92 |
| ***OsHCALDH2/3*-DKO** | 66.6 ± 1.7 | 0.04 ± 0.01 | 76.6 ± 3.0 | 0.19 ± 0.08 | 48.1 ± 2.3 | 0.05 ± 0.02 | 41.9 ± 10.6 | 1.20 ± 0.68 | 33.4 ± 6.4 | 1.65 ± 0.35 | 43.6 ± 10.5 | 0.57 ± 0.23 |
| **Fractionation (%)** | *slow* | *fast* | *slow* | *fast* | *slow* | *fast* | *slow* | *fast* | *slow* | *fast* | *slow* | *fast* |
| **WT** | 91.8 ± 0.8 | 8.2 ± 2.4 | 95.0 ± 0.3 | 5.0 ± 0.6 | 88.5 ± 1.7 | 11.5 ± 5.1 | 77.4 ± 3.2 | 22.6 ± 6.8 | 60.5 ± 3.9 | 39.5 ± 4.5 | 60.7 ± 5.6 | 39.3 ± 5.8 |
| ***OsCAld5H1*-KO** | 88.3 ± 2.2 | 11.7 ± 3.0 | 87.3 ± 1.4 | 12.7 ± 1.5 | 88.4 ± 1.1 | 11.6 ± 2.8 | 77.6 ± 1.9 | 22.4 ± 2.6 | 65.0 ± 3.0 | 35.0 ± 8.8 | 51.8 ± 7.1 | 48.2 ± 7.2 |
| ***OsCAld5H1*-OX** | 90.6 ± 1.0 | 9.4 ± 1.1 | 90.2 ± 1.0 | 9.8 ± 1.1 | 86.1 ± 2.1 | 13.9 ± 4.0 | 75.7 ± 1.3 | 24.3 ± 1.8 | 58.8 ± 2.3 | 41.2 ± 2.7 | 63.0 ± 3.7 | 37.0 ± 3.9 |
| ***OsC3'H1*-KD** | 85.6 ± 1.5 | 14.4 ± 2.6 | 84.8 ± 2.2 | 15.2 ± 3.2 | 74.5 ± 3.2 | 25.5 ± 7.2 | 68.2 ± 4.8 | 31.8 ± 5.1 | 70.5 ± 4.1 | 29.5 ± 5.6 | 71.8 ± 4.4 | 28.2 ± 7.9 |
| ***Os4CL3*-KO** | 81.6 ± 0.7 | 18.4 ± 2.1 | 92.5 ± 3.1 | 7.5 ± 7.0 | 89.6 ± 0.7 | 10.4 ± 0.8 | 88.5 ± 2.7 | 11.5 ± 4.2 | 71.4 ± 3.9 | 28.6 ± 4.3 | 65.6 ± 6.6 | 34.4 ± 6.5 |
| ***Os4CL4*-KO** | 78.5 ± 2.6 | 21.5 ± 6.0 | 75.8 ± 3.0 | 24.2 ± 6.6 | 68.1 ± 1.5 | 31.9 ± 4.5 | 79.1 ± 6.4 | 20.9 ± 9.6 | 55.4 ± 4.1 | 44.6 ± 4.5 | 55.5 ± 6.8 | 44.5 ± 7.2 |
| **WT2** | 93.9 ± 0.8 | 6.1 ± 3.3 | 94.7 ± 0.4 | 5.3 ± 0.9 | 92.5 ± 0.8 | 7.5 ± 1.9 | 73.7 ± 3.4 | 26.3 ± 4.0 | 64.5 ± 4.7 | 35.5 ± 4.9 | 59.8 ± 7.1 | 40.2 ± 7.1 |
| ***OsHCALDH2/3*-DKO** | 81.9 ± 0.5 | 18.1 ± 1.6 | 85.6 ± 1.1 | 14.4 ± 1.9 | 79.6 ± 1.2 | 20.4 ± 3.4 | 66.6 ± 7.3 | 33.4 ± 7.7 | 53.1 ± 4.7 | 46.9 ± 4.8 | 55.6 ± 4.8 | 44.4 ± 5.9 |

Delay time–dependent signal decay data for the major cellulose (**$C_1$**, **$C_{4a}$**, **$C_{4b}$**, **$C_{6a}$**, and **$C_{6b}$**) and lignin methoxy (**OMe**) carbon sites (**Table S2**) were fitted using a double exponential function to determine two independent $T_1$ values for the slower- and faster-relaxing components. The fitted parameters and the standard deviations of the fitting coefficients are listed. WT1 and WT2, wild-type control lines; *OsCAld5H1*-KO, *OsCAld5H1*-knockout line; *OsCAld5H1*-OX, *OsCAld5H1*-overexpressing line; *OsC3'H1*-KD, *OsC3'H1*-knockdown line; *Os4CL3*-KO, *Os4CL3*-knockout line; *Os4CL4*-KO, *Os4CL4*-knockout line; *OsHCALDH2/3*-DKO, *OsHCALDH2*- and *OsHCALDH3*-double-knockout line. The WT2 and *OsHCALDH2/3*-DKO lines, as well as WT1 and all other transgenic lines, were grown side-by-side.

**Table S4. WoodSAS fitting data of dry-state rice cell walls.**

| Genotype | *A* | $2\overline{R}$ (Å) | $\Delta\overline{R}/R$ | a (Å) | *Δa/a* | *B* | σ (× $10^{-2}$) | *C* (× $10^{-3}$) | *α* |
|---|---|---|---|---|---|---|---|---|---|
| **WT1** | 21.0 ± 1.4 | 25.9 ± 0.6 | 0.12 ± 0.01 | 31.3 ± 1.3 | 0.43 ± 0.04 | 17.2 ± 3.0 | 5.3 ± 0.68 | 0.31 ± 0.16 | 4.01 ± 0.04 |
| ***OsCAld5H1*-KO** | 23.1 ± 3.7 | 25.6 ± 0.5 | **0.15 ± 0.01**** | 31.8 ± 1.9 | 0.43 ± 0.06 | 18.8 ± 8.3 | 5.3 ± 0.35 | 0.30 ± 0.06 | 4.04 ± 0.03 |
| ***OsCAld5H1*-OX** | 22.6 ± 1.8 | 25.9 ± 0.1 | 0.13 ± 0.06 | **28.5 ± 0.6**** | 0.51 ± 0.06 | 21.8 ± 2.4 | **2.6 ± 0.39**** | 0.31 ± 0.09 | 4.01 ± 0.03 |
| ***OsC3'H1*-KD** | 20.3 ± 1.9 | 25.3 ± 0.1 | 0.12 ± 0.01 | 32.0 ± 1.4 | **0.35 ± 0.00*** | **32.9 ± 4.4**** | 5.8 ± 0.17 | 0.49 ± 0.19 | 4.01 ± 0.05 |
| ***Os4CL3*-KO** | **18.0 ± 1.4*** | 25.5 ± 0.4 | 0.11 ± 0.03 | **28.8 ± 1.0*** | **0.49 ± 0.01*** | 19.0 ± 1.8 | **2.3 ± 0.29**** | 0.27 ± 0.04 | 4.04 ± 0.01 |
| ***Os4CL4*-KO** | **6.5 ± 1.7**** | 24.9 ± 1.1 | **0.21 ± 0.07*** | **42.0 ± 5.4**** | 0.39 ± 0.08 | 22.8 ± 10.3 | 5.1 ± 1.36 | 0.28 ± 0.07 | 4.04 ± 0.03 |
| **WT2** | 29.0 ± 2.6 | 26.4 ± 0.3 | 0.14 ± 0.02 | 33.3 ± 2.4 | 0.53 ± 0.03 | 22.5 ± 2.9 | 3.6 ± 0.37 | 0.31 ± 0.05 | 4.07 ± 0.01 |
| ***OsHCALDH2/3*-DKO** | 24.8 ± 3.6 | 26.7 ± 0.7 | 0.21 ± 0.06 | 31.8 ± 1.3 | 0.53 ± 0.05 | 20.3 ± 3.3 | 3.4 ± 0.15 | 0.36 ± 0.06 | 4.04 ± 0.06 |

The values are means ± standard deviation of biologically independent rice culm samples ($n$ = 4). Asterisks (*) and numbers in bold indicate a significant difference from wild-type plants (Student's *t*-test, **$p < 0.01$, * $p < 0.05$). WT1 and WT2, wild-type control lines; *OsCAld5H1*-KO, *OsCAld5H1*-knockout line; *OsCAld5H1*-OX, *OsCAld5H1*-overexpressing line; *OsC3'H1*-KD, *OsC3'H1*-knockdown line; *Os4CL3*-KO, *Os4CL3*-knockout line; *Os4CL4*-KO, *Os4CL4*-knockout line; *OsHCALDH2/3*-DKO, *OsHCALDH2-* and *OsHCALDH3*-double-knockout line. The WT2 and *OsHCALDH2/3*-DKO lines, as well as WT1 and all other transgenic lines, were grown side-by-side.
